# High-throughput phenotyping methods for quantifying hair fiber morphology

**DOI:** 10.1101/2020.11.24.392191

**Authors:** Tina Lasisi, Arslan A. Zaidi, Timothy Harding Webster, Nicholas Bradley Stephens, Kendall Routch, Nina Grace Jablonski, Mark David Shriver

**Author notes:** Address all correspondence to: Tina Lasisi, Department of Anthropology, Penn State University.

## Abstract

Quantifying the continuous variation in human scalp hair morphology is of interest to anthropologists, geneticists, dermatologists and forensic scientists, but existing methods for studying hair form are time-consuming and not widely used. Here, we present a high-throughput sample preparation protocol for the imaging of both longitudinal (curvature) and cross-sectional scalp hair morphology. Additionally, we describe and validate a new Python package designed to process longitudinal and cross-sectional hair images, segment them, and provide measurements of interest. Lastly, we apply our methods to an admixed African-European sample (n=140), demonstrating the benefit of quantifying hair morphology over qualitative classification or racial categories, and providing evidence against the long-held belief that cross-sectional morphology predicts curvature.

## Introduction

Human scalp hair morphology is an important, but poorly understood phenotype that varies considerably within and among populations. Scalp hair morphology--alternatively described as hair texture, form, shape or type--refers to the structural appearance of the hair shaft protruding from the follicle. Hair morphology can be examined at multiple scales, from characterization of the cortical cells, medulla, and cuticle to descriptions of overall macroscopic “texture” perceived when considering a head of hair in its entirety. Variations in hair morphology have been linked to variation in DNA sequence, as well as in cellular, protein, and chemical structure^1^. However, focus on investigating the underlying causes of the perceived macroscopic variation has come at the expense of developing language and methodology for describing the phenotype itself. This need is illustrated by the multitude of subjective and, at times, race-based classification systems used across disciplines interested in the variability of this trait^2–5^.

The morphology of an individual hair shaft is the most immediate “macroscopic” scale after considering a head of hair as a whole. At hair-shaft scale, two objectively quantifiable aspects of hair morphology can be delineated: its longitudinal curvature and its cross-sectional geometry (see Fig. 1).

**Figure 1.**
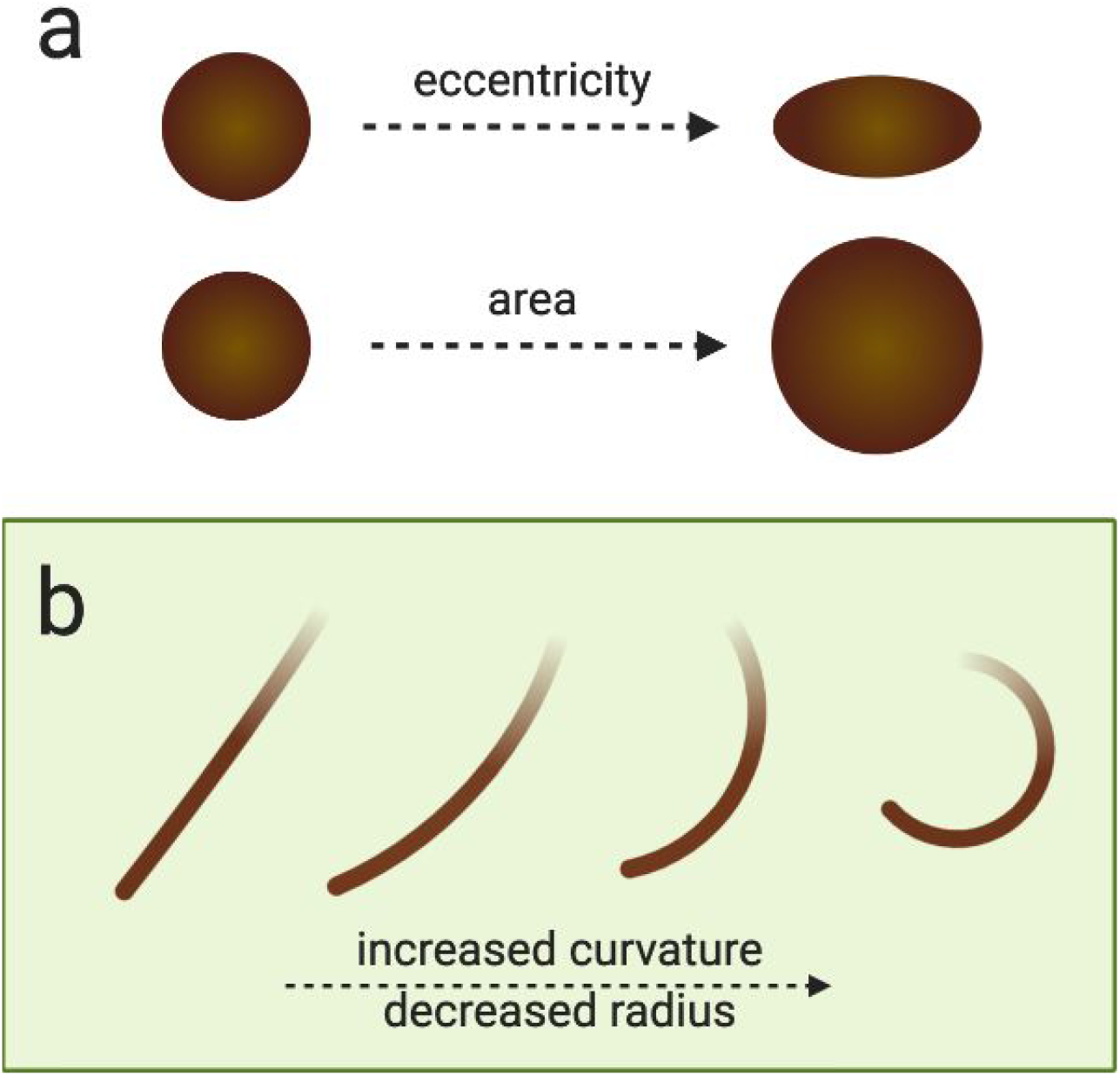
Quantifiable aspects of hair morphology. **a**, Diagram of cross-sectional eccentricity and area. **b**, Curvature is calculated as the inverse of the radius in mm (*radius*^−1^), ergo an increase in curvature corresponds with a decrease in the radius of the circle fitting the curve.

Work in this field has been plagued by a lack of standardization in methods and issues of replicability, in part due to inadequate detailing of methods used and subjectivity in their application ([Supplementary Information](https://drive.google.com/open?id=1bIwpFyeX2_RvYYk92kJ4kW2fG4d3xD8OoWHeIe_h3Kc)). In light of these challenges, we have developed sample preparation and image analysis methods that allow for the high-throughput phenotyping of hair fiber cross-sectional geometry and curvature. Our aim was to develop methods that would 1) be appropriate for the full range of human hair diversity, 2) minimize or eliminate subjective observer input, 3) require no specialized skills or equipment, and finally, 4) be efficient and scalable. Here, we present a comprehensive description of the protocols used for sample preparation and a novel computational tool for the analysis of images created with those protocols.

## Results

### Using a low melt point plastic for immediate embedding of multiple hairs

Cutting hairs at a perpendicular angle is crucial for the accurate visualization of a fiber’s cross-section. Traditional methods using resin or paraffin require long (~24h) curing times and make it difficult to embed curled hairs for sectioning. We found that a low melt point plastic such as polycaprolactone allowed us to lay multiple hairs of any morphology in parallel lines. The material immediately encases curled hairs even when they are stretched straight. For our purposes, we embedded six hairs from each individual using polycaprolactone plastic sheets (Polly Plastics, Michigan, USA). We heated a strip of moldable plastic on a hot plate and stretched hair samples over this strip affixing them to the material. We then placed a second strip of plastic on top of the hairs and put a heated block on top to fuse both strips of plastic and embed the hairs completely. After storing the embedded samples in a 4C room for a minimum of two hours and we sectioned them with a PanaVise 507 Flat Ribbon Cable Cutter fitted to a PanaVise 502 Precision PanaPress. We found that a regular razor blade was equally capable of cutting through the samples, but the PanaVise set up allowed us to process samples at a higher rate. We then mounted the sectioned samples upright between two Plexiglas blocks for visualization and imaged with a Leica DMLS microscope (Leica, Wetzlar, Germany) at 10x with a Lumina GX8 camera (Panasonic, Osaka, Japan) attachment. Illumination was provided through the sides of the Plexiglas hair chip support using fiber optic dissecting scope lights See Figure 2a, [Supplementary Video 1](https://youtu.be/OSgiErHmiDE), and Methods for step-by-step protocol.

**Figure 2.**
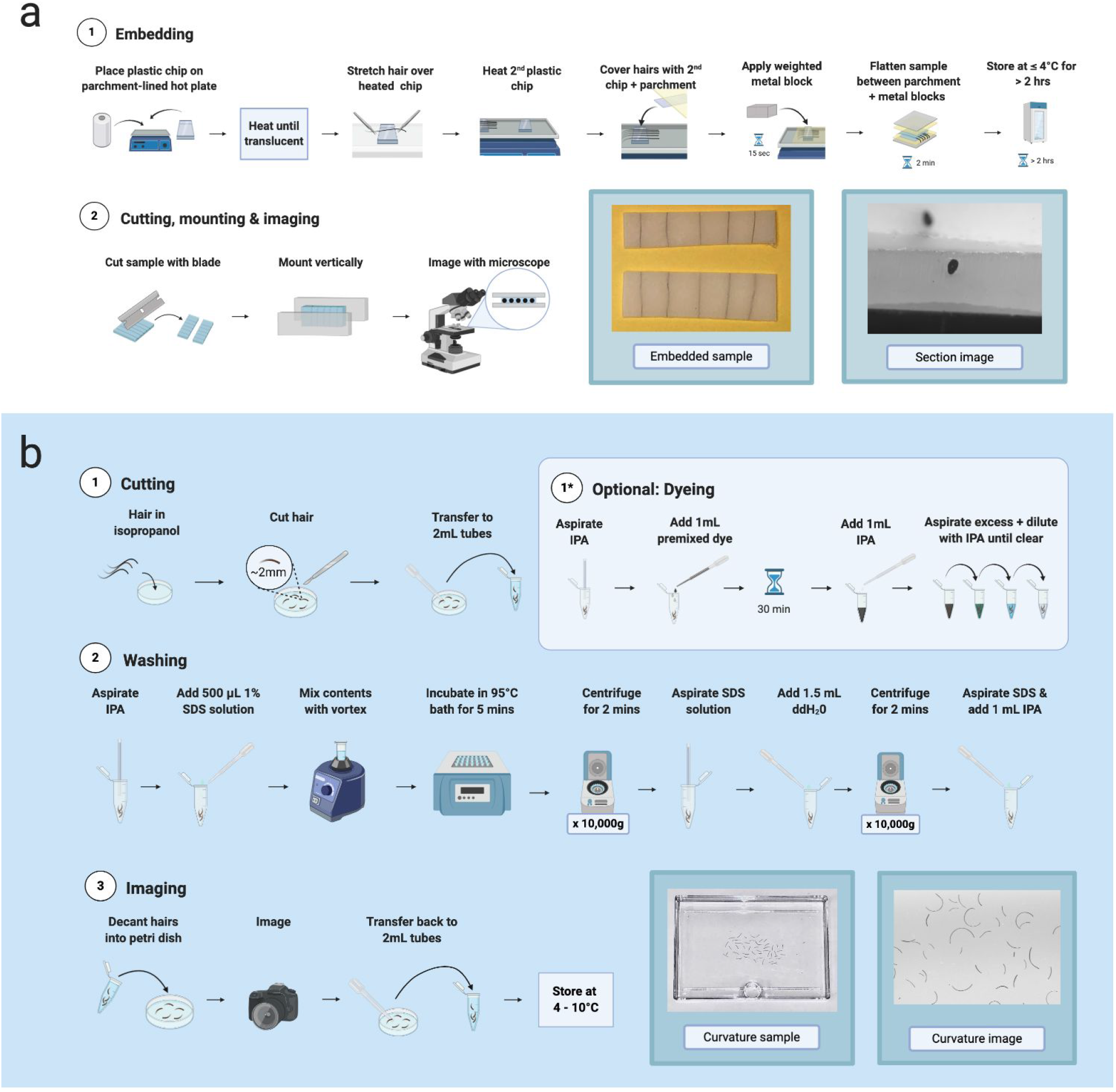
Flowcharts of laboratory protocols used. **a**, Embedding protocol for cross-sectional imaging. **b**, Sample preparation protocol for curvature imaging.

### Cutting hair fibers into fragments as a scalable method for washing and imaging hair fibers for curvature analysis

Multiple factors associated with grooming can temporarily alter the curvature of a hair fiber (e.g. hair products, straightening irons, braiding). The first step of our sample preparation was developed to remove the effect of these extrinsic factors and to allow for the measurement of curvature in two dimensions. To achieve this, we cut hairs into small fragments and used a multi-step washing process to remove any residue and allow the hairs to revert to a shape representing the fiber’s intrinsic curvature. We used three to five hair fibers from the crown of each individual in order to capture a representative value of hair curvature for that individual. We placed hairs into a Petri dish containing 5mL of isopropyl alcohol (IPA) and cut them into fragments of 3mm with a curved-point scalpel blade. We then transferred them to 2mL tubes using transfer pipettes. At this stage, lightly pigmented hairs were dyed black using a commercial hair dye kit to improve final imaging contrast. We washed hairs with a sodium dodecyl sulfate (SDS) mixture (1% SDS, 99% double distilled H2O) and rinsed them with H2O. Finally, we stored hairs in 1mL of IPA until imaged. For imaging, we decanted each sample into a petri dish containing 5mL of IPA and we used a Panasonic GH4 camera with Olympus f2.8 60mm macro lens to capture the images (see Fig. 2b, [Supplementary Video 2](https://youtu.be/qR-Jj4K0Bts), and Methods for step-by-step protocol).

### Automated image analysis for unsupervised high-throughput processing

Our Python package, *fibermorph*, is a user-friendly, fully automated image analysis tool that provides a convenient way to estimate hair eccentricity and curvature from cross-sectional and longitudinal images (see Fig. 3). We designed this package to run on the command line and provide detailed guidance on its use so that any researchers who have images of hair sections or curvature can use it without programming experience. No comparable tools exist for these purposes, nor are there any computational tools that are consistently employed by researchers studying hair fiber curvature and cross-sectional morphology.

**Figure 3.**
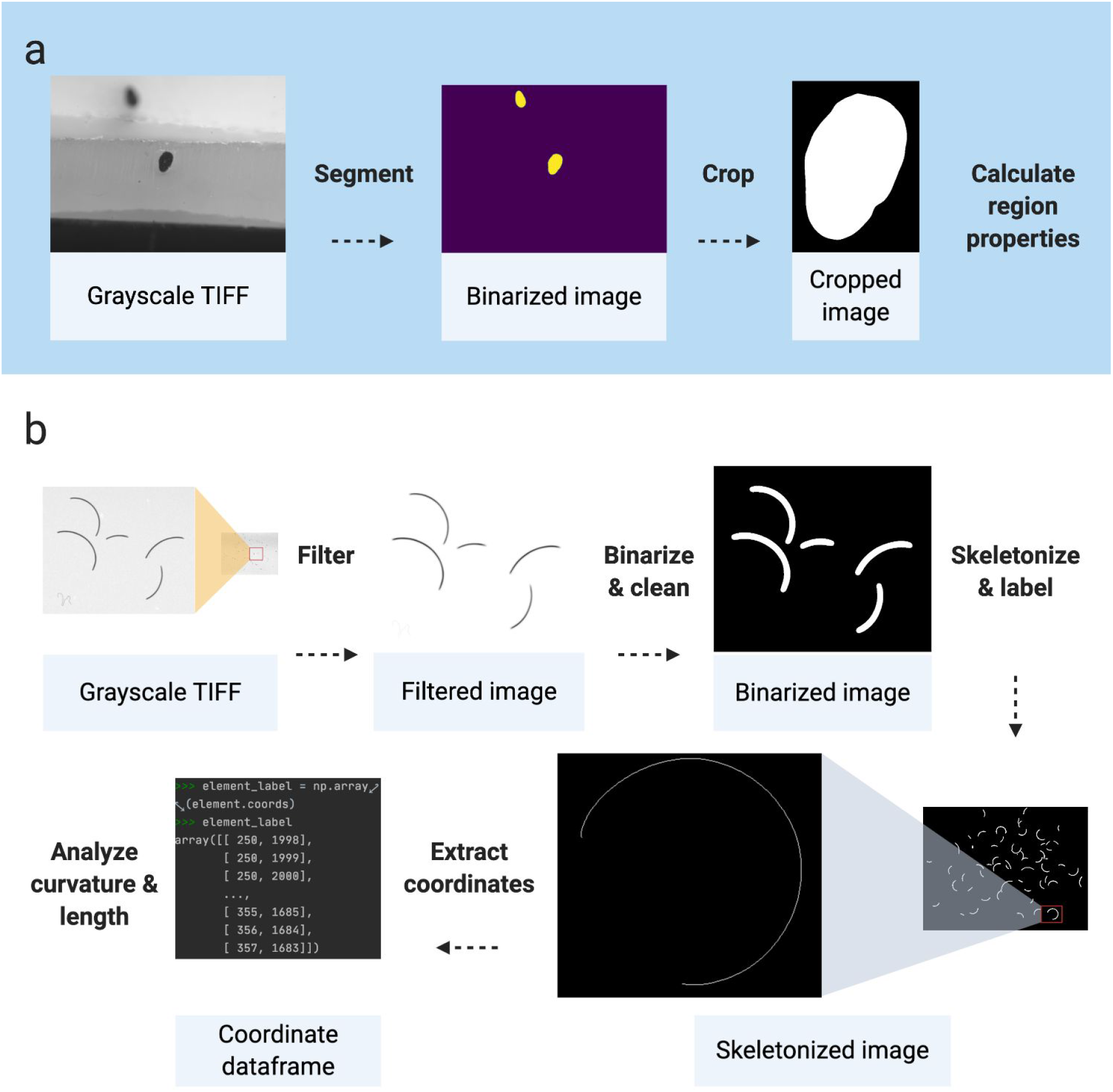
fibermorph image analysis workflow. **a**, Cross-sectional image processing and analysis. **b**, Curvature image processing and analysis.

### Measurement error in hair fiber curvature estimation

*fibermorph*, is designed to simultaneously estimate curvature of multiple hair fragments in an image. The program first processes the image to extract hair fibers from the background and reduces each hair to 1 pixel width. The pixel coordinates are then used to estimate hair curvature using a circle fitting algorithm, returning the curvature for each fragment measured as the inverse of the (fitted) circle’s radius. The program returns a spreadsheet for each image containing the image ID, mean and median curvature (across all hair fragments in the image), hair count, and mean and median length.

To test the accuracy of *fibermorph’s* curvature estimation, we simulated 20 images with a range of curvatures. Each image contained 25 hair fragments of the same length and curvature, but different orientations, representing a sample of hair collected from a single individual. The simulated curvature ranged from 0.1mm^−1^ to 2mm^−1^ representing the observed range of curvature in our sample of real hair. We used *fibermorph* to estimate the mean and median curvature and length of hair in each image and compared it to the known (simulated) length and curvature (see Fig. 4).

**Figure 4.**
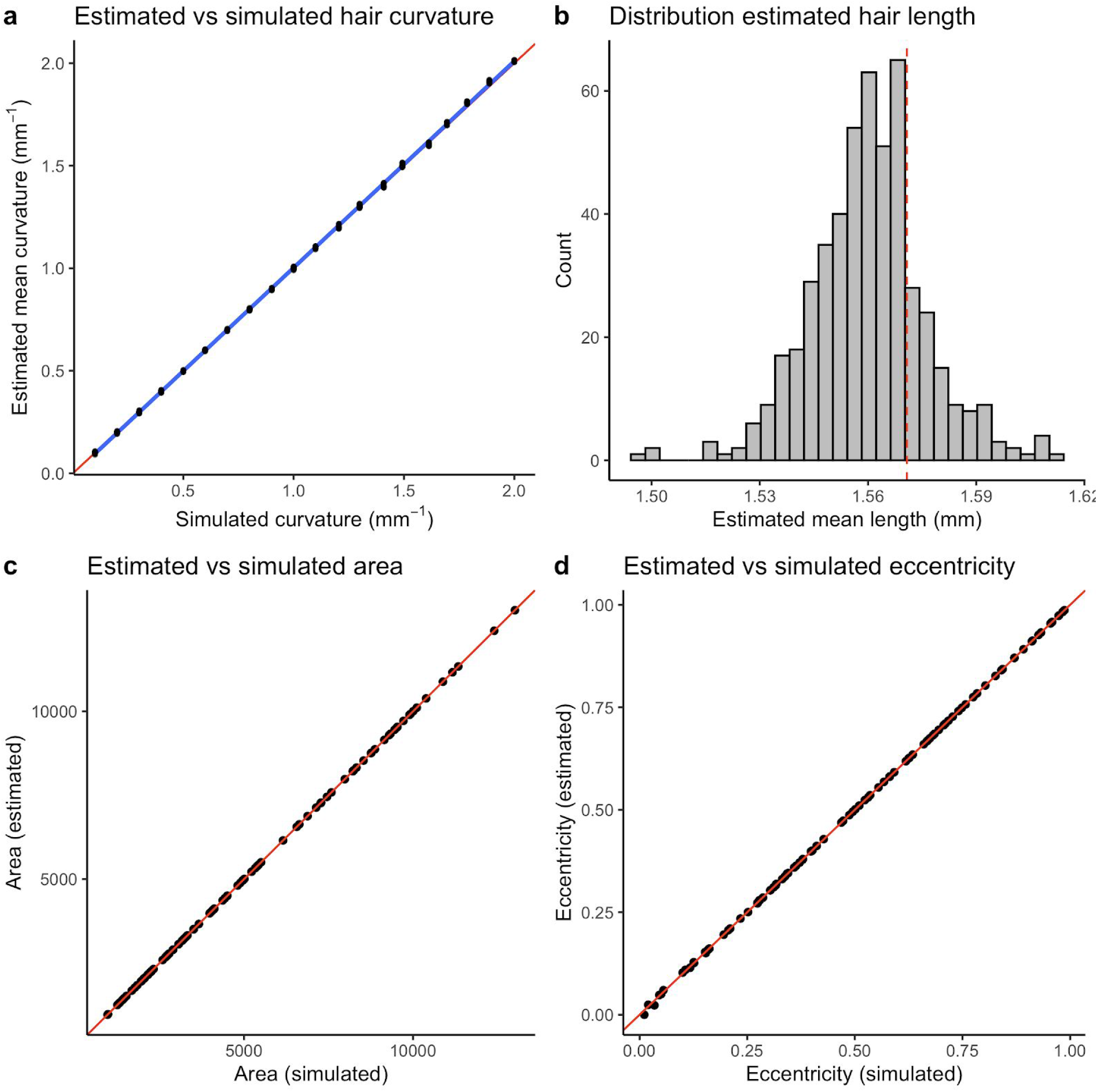
Performance of *fibermorph* on simulated data. **a,** Correlation between estimated and true curvature. The red line represents y = x and the blue line represents the line of best fit. **b,** Distribution of hair fragment length estimates. The dashed red line shows the true simulated length of each fragment. **c,** Correlation between estimated and true cross-sectional area. **d,** Correlation between estimated and true eccentricity.

*fibermorph* accurately estimates hair curvature across a range of simulated curvature values (r^2^ = 0.999; Fig. 4a). The error in estimation was minimal overall with mean root mean squared error (RMSE) of 2.21 × 10^−4^ mm^−1^ (0.47%) The estimated length of each hair fragment was similarly accurate (Fig. 4b) with an RMSE of 4.31 × 10^−4^ mm (0.69%). See Supplementary Results 1 for details on RMSE and percent error for curvature.

### Measurement error in cross-sectional parameter estimation

*fibermorph* also processes micrographs of hair sections and measures a number of cross-sectional parameters. The program first crops the images, then segments out the cross-section from the background using scikit-image’s implementation of the Chan-Vese algorithm^6,7^. The binarized image is then used to calculate area, minimum diameter, maximum diameter in microns (μm), and eccentricity, which is a measure of how elliptical the cross-section is (see Methods for details).

To evaluate the measurement error in estimation of cross-sectional parameters, we simulated 100 ellipses with minimum and maximum diameters chosen at uniform intervals between 30 to 120 μm. The range of minimum and maximum diameters is based on the range found in human scalp hairs. fibermorph accurately estimates the area and eccentricity of ellipses (Fig. 4c). The RMSE for area is 0.51 μm^2^ (0.01%) and 6.49 × 10^−4^ (1%) for eccentricity. See Supplementary Results 1 for details on RMSE and percent error for cross-sectional geometry.

### Self-reported hair texture and “objective” classification fail to capture quantitative variation in curvature

We compared quantitative hair fiber curvature with self-reported hair texture in a sample of 140 individuals of admixed European and African ancestry. Self-reported hair texture or form is often used for phenotyping purposes in genome-wide association studies (GWAS) with ordinal categories such as “straight”, “wavy” and “curly”. However, we find that while there is a correlation between these ordinal categories, individuals are inconsistent in their perception of hair texture (see Fig. 5a). In other words, there seems to be variation in the level of perceived hair curl each of these categories encompass. An alternative to subjective hair form categorization is the classification of hair into ordinal categories based on their objective curvature ^5^. We also analyzed the variation in hair curvature using the curvature thresholds described in the 2007 Loussouarn et al. paper ^5^ (see Fig. 5b) and found that the continuous variation was binned in a manner that unequally represented variation across the categories.

**Figure 5.**
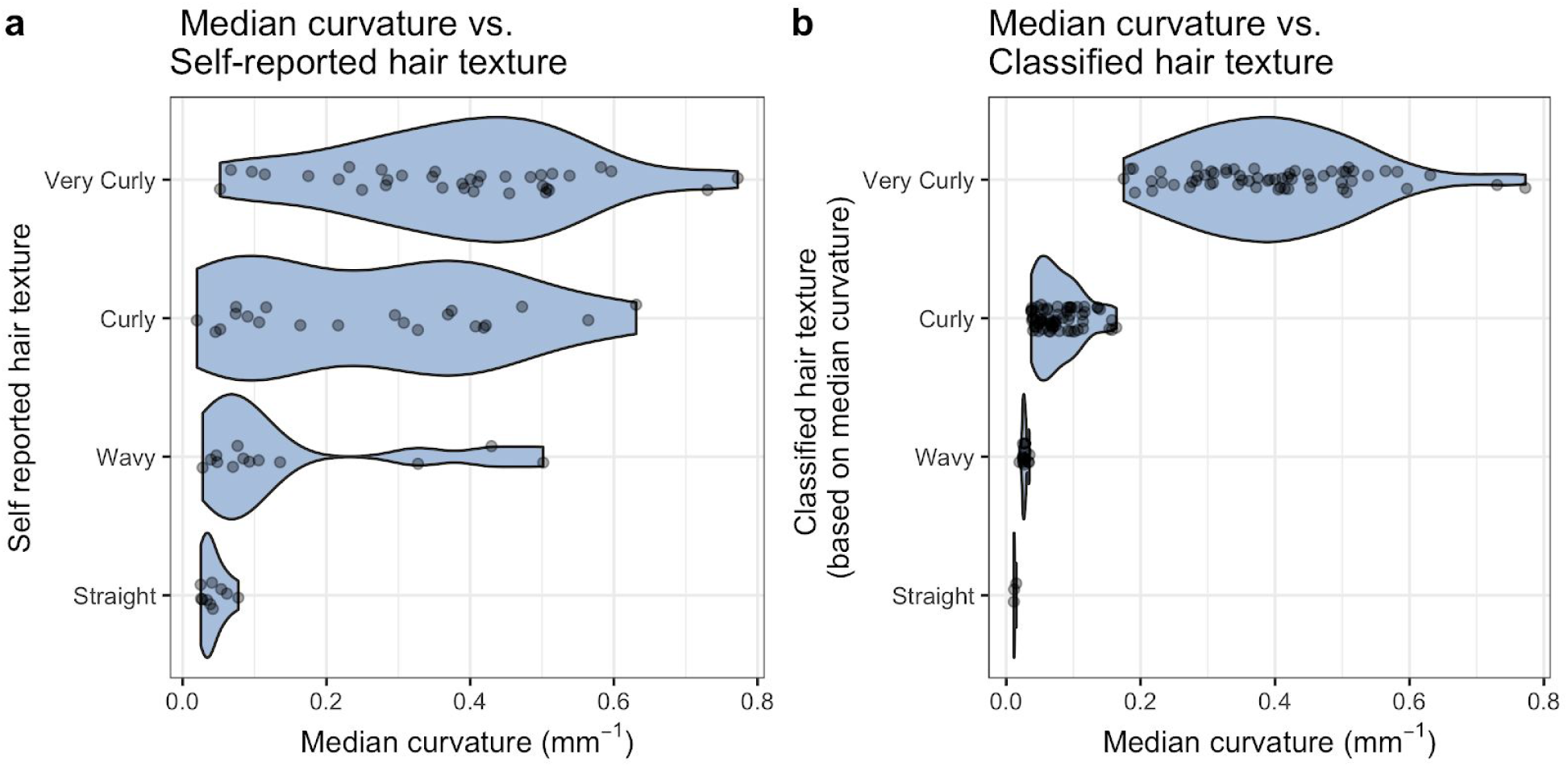
Comparison of quantitative and qualitative hair morphology. a, Self-reported hair form shows inconsistency among individuals with regards to the level of curvature encompassed by each category. **b**, Classifying hair based on objective curvature Loussouarn et al. 2007, obscures the wide range of variation that exists in the “Very Curly” category.

### Quantitative hair fiber morphology elucidates the relationship between curvature and cross-sectional shape and genetic architecture

Many studies have reported a correlation between hair fiber curvature and cross-sectional shape, an observation which has been interpreted as a causal effect of eccentricity on hair curl. Specifically, the presence of elliptical cross-sections in populations with tightly curled hair (i.e. West African) and the presence of round cross-sections in populations with straight hair (i.e. East Asian) has been interpreted as evidence that cross-sectional shape of hair dictates its curvature^8,9^. Whether these two traits are genetically and/or developmentally correlated is unknown because of population structure, which induces spurious correlations between traits^10^. For example, individuals with more West African ancestry tend to have more pigmented skin and curlier hair, on average, than individuals with more European ancestry. However, there is no reason to believe that these two traits are correlated because of a shared genetic architecture. We illustrate this in a sample of 140 individuals with mixed African and European ancestry. We show that the positive association between melanin index and hair curvature (slope t-statistic = 12.50, p-value < 2 × 10^−16^, Fig. 6c) is no longer significant when proportion of African ancestry is used as a covariate in the model to correct for ancestry stratification (slope t-statistic = 1.86, p-value = 0.065, Fig. 6d). Similarly, we show that the observed positive relationship between eccentricity and hair curvature (slope t-statistic = 6.08, p-value = 1.96 × 10^−8^, Fig. 6a) is also not significant when corrected for proportion of African ancestry (slope t-statistic = 0.87, p-value = 0.386, Fig. 6b). This demonstrates that the correlation between hair curvature and cross-sectional shape in people of mixed African and European ancestry, as well as between Africans and Europeans, is driven by population structure, similar to the correlation between melanin index and curvature. More detailed analyses (e.g. estimation of genetic correlation and GWAS of both traits) in a larger sample will be needed to elucidate whether some component of the relationship between hair curvature and cross-sectional shape results from pleiotropy. See Supplementary Results 2 for full set of analyses and admixture breakdown of the sample.

**Figure 6.**
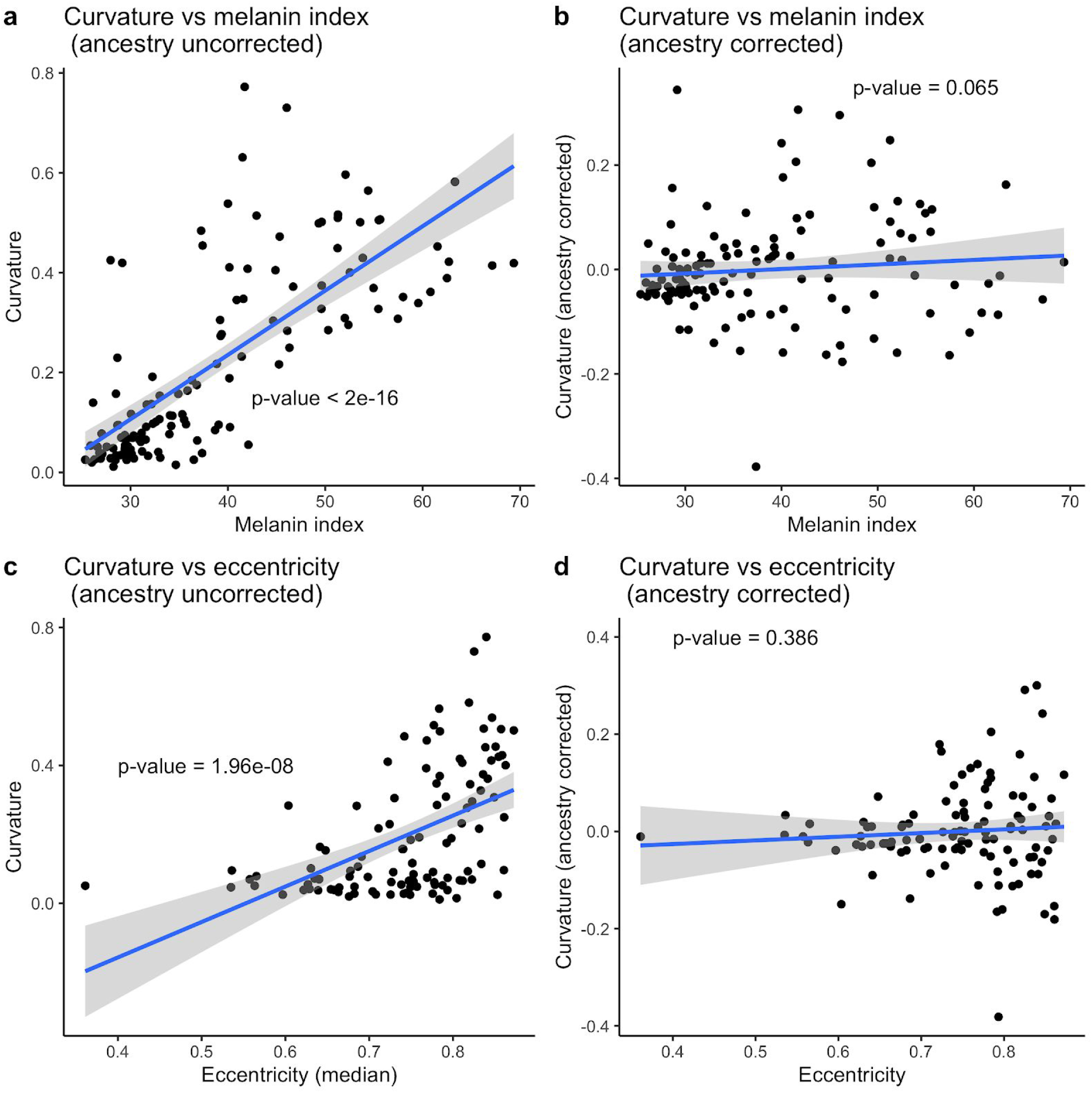
The effect of population structure on trait-trait correlations. **a**, The correlation between hair curvature and melanin index without controlling for ancestry. **b**, The correlation between hair curvature and melanin index controlling for African ancestry. **c**, The correlation between hair curvature and cross-sectional eccentricity without controlling for ancestry. **d,** The correlation between hair curvature and cross-sectional eccentricity controlling for African ancestry.

## Discussion

Our results demonstrate that the methods we have developed can reliably quantify cross-sectional morphology and curvature in hair fibers. Moreover, we illustrate that the practice of categorizing hair either subjectively or by using arbitrary thresholds mischaracterizes the distribution of quantitative hair morphology, specifically, underestimating the variation that exists in more tightly curled hair.

Our sample preparation protocols and computational image analysis tool represent significant methodological advancements over previous attempts to quantify hair morphology. The sample preparation protocol for curvature imaging described here is considerably more scalable than methods used in other studies of hair curvature. Our protocol introduces a standardized method of washing hair samples that more closely resembles protocols used in sample preparation for trace element analysis in hair^11,12^ and that is superior to previous methods that required handling of individual hairs with forceps^13,14^. Our method operationalizes the measurement of curvature by defining intrinsic curvature as the curvature that a hair fiber has when reduced to a short length (~1-3mm) which reduces the three dimensional curvature of certain hairs to a two dimensional curve that can be measured without the distortion that was needed in previous methods or the weight of the hair fiber itself.

Our embedding protocol using the low melt-point plastic (polycaprolactone) overcomes the problems of traditional embedding techniques that used resin or paraffin^9,13^. These techniques are feasible for straight hairs, which can be easily manipulated to lie flat and parallel to each other, but are not practical for curly hairs because non-straight hairs cannot be reliably embedded in a manner that allows for a reproducible cross-sectional cut. Previous attempts to overcome this problem included using a heat-shrink tube and drawing the hair through the tube^15^, bundling hairs and slightly embedding them before embedding them fully^16^, and stretching hairs over cardboard before embedding them in a large resin block^13^. All of these methods are laborious and cannot be scaled to the study of large samples. Our method allows for the immediate embedding of hairs, regardless of shape, allowing researchers to prepare dozens of samples per hour.

Our novel computational tool streamlines the analysis of curvature and cross-sectional geometry of hair fibers. This tool requires no input from a user (other than the location of the image files), and so removes inter-observer error and subjectivity in assessing curvature^13,17^. It also saves time and improves accuracy and reproducibility because tasks that would have previously required hours of labor can now be executed unsupervised by an automated program that requires no additional cost or training to use. In the absence of comparable computational tools for hair fiber morphology, we demonstrated, in detail, the technical validity of these methods by testing *fibermorph’s* performance on simulated geometric shapes of known parameters. Most importantly, the laboratory protocols are hosted in open-access repositories that allow for easy feedback and modification, and the code for the image analysis has been made open source to facilitate collaboration and further elaboration.

Our results show that both self-reported hair texture and “objective” classification obscure a considerable amount of variation (see Figure 5b). Moreover, our analysis of quantitative curvature compared to self-reported hair texture suggests that participant interpretations of subjective curl categories are inconsistent (see Fig. 5a). By applying “racial” categorization to hair, forensic scientists and dermatologists alike are bound in tautology that results from racial stereotyping, i.e. “African/Asian/European” individuals have “African/Asian/European” hair morphology and vice versa, the hair of a particular “racial” morphology is found only in those “races”^18–20^. This presents a paradigm wherein individuals, and entire populations, who fall outside of those options cannot be considered or characterized. By applying our methods to an admixed African-European sample, we demonstrate the potential for uncovering the genetic architecture of hair morphology (see Figure 6b).

Researchers interested in understanding the biology underlying variation in hair fiber curvature are severely hindered by the oversimplification resulting from the use of qualitative hair types because these typologies preclude the analysis of the features that may independently contribute to macromorphology (e.g. the effect of cross-sectional eccentricity on curvature). In this sample, we were able to show that eccentricity does not predict hair curvature, which is in line with the findings of other studies^8,13,21^. In fact, we explain this correlation as arising due to uncorrected population structure, which induces correlations between traits that are not necessarily genetically or physically linked. Nevertheless, the persistence of this idea demonstrates the importance of using quantitative methods to disentangle the factors contributing to hair morphology rather than relying on descriptive comparisons between racialized groups or ethnicities.

Human scalp hair morphology is a complex phenotype that has remained poorly understood due to the high threshold of investment in time and resources required to apply existing quantitative methods. With these high-throughput phenotyping methods, we provide researchers with a comprehensive starting point and the option to identify specific components to focus on befitting their specializations. Most importantly, due to the open access and open source infrastructure of this project, collaboration is facilitated and democratized, allowing anyone who is interested to work on this research. This work represents a much needed baseline and standardization that is fundamental to the incremental improvements that were previously unfeasible in the study of this complex phenotype.

## Methods

### Hair samples and genotype data

The data consists of 140 hair samples and corresponding genotype data from individuals of admixed European and African ancestry.^22^ These were collected as part of a larger study (Anthropometry, DNA, and Perception of Traits or ADAPT) with informed consent and ethical approval by the Pennsylvania State University Institutional Review Board (#44929 and #45727). To select these individuals, we merged the full genotype dataset (n=4257 individuals genotyped on the 23andMe V4 array) with the 1000 Genomes reference panel.^23^ We pruned SNPs for linkage disequilibrium (PLINK 1.9 “--indep-pairwise 100 10 0.” 1 yielding a set of 118x SNPs) and estimated genomic ancestry using an unsupervised clustering approach (k=5) with ADMIXTURE.^24,25^ We selected individuals with >80% combined African and European ancestry and <10% ancestry from any other group for whom hair samples (more than 4 hair fragments per person) and skin reflectance were available (n=140). The hair samples and genotype data for the admixed individuals were collected with informed consent and ethical approval by The Pennsylvania State University Institutional Review Board (#44929 and #45727).

### Hair embedding, sectioning and imaging protocol

In using polycaprolactone plastic sheets, we found that embedding hairs in this low melting point plastic offered considerable benefits compared to the use of resin. We were able to immediately encase hairs in the plastic, which solved the challenge of keeping curled hairs stretched in position for 24 hours when embedding in resin. For our purposes, we used Polly Plastics (Michigan, USA) moldable plastic sheets which are readily available and affordable.

For the embedding process, we cut down the plastic sheets to ~15mm by ~30mm and heated them on a hot plate (lined with parchment paper) for 15 seconds until translucent and softened. For each hair sample, we embedded six hairs by stretching hairs in parallel lines over the heated plastic strip to encase them in the plastic. A second strip of heated plastic was then placed over the strip containing the embedded hairs. A heated steel block was then placed on top of the sample for 15 seconds to fuse both strips of plastic and completely embed the hairs. The sample was then removed and cooled for 5 minutes before cutting down excess plastic and hairs using a template. The sample was then stored in a 4°C refrigerated room for at least two hours to ensure the plastic was hardened. See [Supplementary Video 1](https://youtu.be/OSgiErHmiDE).

To section the hairs, we were able to use a flat-edged razor, but to facilitate the processing of a high volume of samples, we used a mechanical press with attached razor (PanaVise 502 Precision PanaPress with PanaVise 507 Flat Ribbon Cable Cutter). The low melting point of the plastic means that it will melt at body temperature, so it is important to handle the samples minimally and to section quickly, hence our decision to use this set up over manual sectioning with a razor. Sectioned samples were mounted upright between two clear blocks for visualization and imaged with a Leica DMLS microscope (Leica, Wetzlar, Germany) at 10x with a Lumina GX8 camera (Panasonic, Osaka, Japan) attachment. See step-by-step protocol on Protocols.io.^26^

### Hair sample preparation and imaging protocol for curvature

To obtain a representative sample for each individual we recommend that a minimum of five hair fibers (≤ 30mm) are used per individual. For shorter hairs, a minimum of 10 hairs (≤ 10mm) is recommended. Because hair samples were collected prior to the development of this protocol, we did not have enough hairs for each individual to adhere to this rule. As such, we demonstrate the results of our analyses (described below) with and without low hair count samples.

The first step of our protocol is to cut the hair sample into fragments in a Petri Dish containing 5mL of isopropyl alcohol (IPA). We found that hairs were easier to handle in this medium than in water, where hairs variably sunk to the bottom of the dish or stuck to the tools we used to handle and cut them. Hairs were placed into a Petri dish containing 5mL of isopropanol (IPA) and cut into fragments of 3mm with a new #10 scalpel blade for each sample (these blades were cleaned between the processing of separate batches of samples). We found that scalpels with a curved point were easiest to use in the Petri dish, so other similar scalpel blades (e.g. #20, #21 or #22) could be used depending on preference and availability. Once cut, hairs were transferred into 2mL tubes using a fresh transfer pipette for each sample. Hair samples that appeared lightly pigmented at this stage were dyed using a commercial hair dye kit (L’Oreal Paris Feria 6.3fl oz in the color Bright Black) to ensure adequate contrast in the imaging stage. After cutting (and dyeing, if applicable) hairs were washed with a sodium dodecyl sulfate (SDS) mixture (1% SDS, 99% double distilled H2O) and rinsed with H2O. In this process, we attached tubing to a faucet to create an aspiration system where the flowing water created a vacuum. Using a pasteur pipette attached to the vacuum end of the tubing, we aspirated as much IPA from the samples in the 2mL tubes as we could without disturbing the hairs. We then filled the tubes with the SDS mixture until they reached the 2mL mark. Samples were then mixed using a Vortex mixer and placed in a warm water bath (99°C) for 5 minutes. We found that microcentrifuge tube caps were necessary to securely seal the tubes when in the water bath. The purpose of the warm water bath is to thoroughly clean the hair fibers and to ensure all hairs go through the same process of hydration and dehydration as variability in porosity may affect their morphology. Once removed from the water bath and cooled, the hairs were microcentrifuged to collect all hair fragments at the bottom of the tube. The SDS mixture was then aspirated from the tubes, and we refilled the tubes with H2O and microcentrifuged again to rinse the hairs. Finally, the H2O was aspirated and replaced with 1mL IPA. After this step, hairs were stored in these 2mL tubes until imaged. See [Supplementary Video 2](https://youtu.be/qR-Jj4K0Bts).

To ensure the same resolution for each image, we mounted a camera to a stand and set the focus to the same standard for each sample. Immediately prior to imaging, samples were decanted from their 2mL tubes into a Petri dish containing 5mL of IPA. We found IPA to be a suitable medium as hairs invariably sunk to the bottom of the dish for each sample, ensuring consistency between images and removing the effect of shadows. Additionally, 5mL of IPA was enough to cover the hairs eliminating any glare from surface tension related to hairs protruding through the liquid. We imaged the samples using a Panasonic GH4 camera with Olympus f2.8 60mm macro lens producing images that were 5200 by 3900 pixels and at a resolution of 132 pixels per mm. A complete step-by-step protocol with images and video can be found on Protocols.io.^27^

### Image analysis protocol for cross-section

The fibermorph section analysis program requires grayscale TIFF images as input. Where necessary, images are cropped as part of the preprocessing pipeline. Then, images are segmented using a Chan-Vese algorithm^6^ in scikit-image^7^. This algorithm was chosen for its ability to segment images with poorly defined edges but significant differences in grayscale intensity between the region of interest and the rest of the image.

The parameters of the section identified are calculated using scikit-image’s *regionprops* function. We output the following: minimum diameter, maximum diameter, area and eccentricity. Eccentricity is defined as:

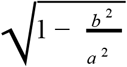

where *b* is the minimum radius and *a* is the maximum radius of the ellipse. Diagrams for all the protocols were created using BioRender.com.

### Image analysis protocol for curvature

The analysis of curvature begins with a grayscale TIFF image file as input. Our pipeline applies a ridge filter to extract the regions of interest (hairs), then binarizes and cleans the image before skeletonizing which reduces each hair fragment to 1 pixel width for the curvature analysis. The functions applied in this process are from the Python library scikit-image. ^7^ The analysis of the processed image starts by labeling each element (hair) and calculating curvature for each of these elements using a Python function based on Taubin's circle fitting algorithm.^28^ The spreadsheet for each sample provides the length and curvature for each fragment and the summary spreadsheet provides the average length and curvature, as well as the number of hairs per image.

The spreadsheets containing the curvature and length measurement for each hair within an image are saved in the analysis folder. The default is to only produce this summary spreadsheet but a user can use a simple command to create a folder with the raw measurements per fragment should they so wish. The final summary spreadsheet contains the mean and medians for the curvatures calculated from those data frames, as well as mean and median length. Length for an element/hair is calculated from the total number of pixels in the thinned element. As the length of horizontal/vertical vs. diagonal pixels is a known issue in image analysis, we implement a correction by counting the number of diagonal pixels using a correction factor of 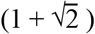 as has been suggested elsewhere.^29^

### Data simulation for curvature and cross-sectional image analysis validation

To test the accuracy of *fibermorph’s* curvature estimation, we simulated 20 images, where each image contained 25 hair fragments of the same length and curvature, but different orientations, representing a sample of hair collected from a single individual. The curvature for each image was chosen with a range of 0.05 to 2 mm^−1^ representing the observed range of curvature in our sample of real hair. To generate randomly oriented arcs, we sampled the start angle (*θ*_*start*_) of each arc from a uniform distribution on the interval (0, *π*) and drew a line through 25 points with angles (*θ*_*i*_) uniformly spaced between *θ*_*start*_ and 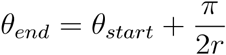 where *r* is the simulated radius. The x and y coordinates of these points were calculated as *x* = *r* × *cos*(*θ*_*i*_) and *y* = *r* × *sin*(*θ*_*i*_).

Our Python script for ellipse simulation creates a canvas of 5200 × 3900 pixels, sets the resolution to 4.25 pixels/μm and uses the chosen minimum and maximum diameters to draw the ellipse using scikit-image. The maximum diameter is chosen first (50 - 120μm), then the eccentricity is chosen from a uniform distribution (0 - 1) and finally, the ellipse is set on an angle from a value chosen from a random distribution (0 - 360°). The parameters (minimum diameter, maximum diameter, area, eccentricity) are saved in a reference spreadsheet and the image is saved as a TIFF with the same name. We simulated 100 ellipses using these parameters for our analyses.^22^

### Analyses with simulated data and real data

We estimated all curvature, length and cross-sectional parameters on both simulated and real data using our *fibermorph* Python package. In our analyses, we used the following R packages: workflowr,^30^ tidyverse,^31^ knitr,^32^ and cowplot^33^

For simulated data, we estimated RMSE as 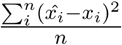 where *x*_*i*_and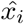 are the true (simulated) and estimated values, respectively. To test whether the correlation between two traits was due to population structure, we fit the following linear model: *y*_*i*_ = *α*+*β*_0_*x*_*i*_+*β*_1_*z*_*i*_ where *y*_*i*_ is the value for one of the traits (e.g. hair curvature), *x*_*i*_ is the value for the other trait (e.g. eccentricity) and *z*_*i*_ is the proportion of African ancestry of the *i*^*th*^ individual. Because our sample is composed of admixed individuals of primarily African and European ancestry, the inclusion of African ancestry as a covariate should correct for the effects of ancestry stratification in our sample^10^. To visualize the correlation between two traits after ancestry correction (e.g. in Fig. 6), we fitted a linear model between the first trait (*y*_*i*_) and African ancestry (*z*_*i*_) and plotted the residuals against the second the trait (*x*_*i*_).

## Ethical approval and informed consent

The hair samples and genotype data for the admixed individuals were collected with informed consent and ethical approval by The Pennsylvania State University Institutional Review Board (#44929 and #45727).

## Supporting information

Supplementary Results 1

Supplementary Results 2

## Data availability

The spreadsheets upset in the analyses are available on Github at https://github.com/tinalasisi/2020_HairPheno_manuscript and the raw images (both simulated and real data) can be found on Zenodo (doi:10.5281/zenodo.4289252).

## Code availability

The code used for the image analysis can be found on GitHub at https://github.com/tinalasisi/fibermorph and the code used for the data analysis can be found at https://github.com/tinalasisi/2020_HairPheno_manuscript.

## Acknowledgements

This research is supported by The Wenner-Gren Foundation (Gr. 9911), the National Science Foundation (No. 1847845) and the Pennsylvania State University Africana Research Center. We would like to thank Alan Rogers and the University of Utah Primate Evolution & Genomics Lab for helpful comments and discussion on earlier drafts of the paper. This development of these methods would not have been possible without the participants recruited for the study and the members of the Shriver Lab (past and present) who helped collect and process this data.

T.L. thanks Jinguo Huang and Tomás González Zarzar for their help in testing out the software. We are especially grateful to Sarita Greer for her supervision of the lab and the undergraduate students involved in parts of the research. We would like to explicitly acknowledge the contribution of the following students in the extensive trial-and-error leading up to the finalization of these methods: Duneshka Cruz, Samantha Muller, Alexis Rollins, Yesenia Hill, Emily Bramel, and Feihong Rodell.

## Author Contributions Statement

T.L. conceived of the methods and directed the project. The experiments with various embedding materials were carried out by K.R., M.D.S. and T.L. The software was written by T.L., N.B.S., and T.H.W. The statistical analyses and relevant code was written by A.Z. and T.L. The manuscript was written by T.L. with input from all the authors. N.G.J. and M.D.S. supervised T.L. during the development of the methods and provided critical feedback in shaping the overall project.

## Notes

https://github.com/tinalasisi/2020_HairPheno_manuscript

https://github.com/tinalasisi/fibermorph

https://pypi.org/project/fibermorph/

### Competing Interest Statement

The authors have declared no competing interest.

