## Supplementary Information for "High-throughput phenotyping methods for quantifying hair fiber morphology"

### Supplementary Information I

### Previous work on cross-sectional geometry

The cross-sectional geometry of human scalp hair has been described in studies dating back to the 19th century(Pruner-Bey, 1864), but the methods for preparing samples for cross-sectional imaging are neither universally applied, nor universally applicable. To observe the cross-section of a hair, it must be cut perpendicularly to its longitudinal axis and magnified with a microscope. Typically, hairs are embedded in an epoxy resin, though examples of other embedding-media can be found (mainly paraffin, which is commonly used in histology). Whereas the width of a scalp hair fiber is on the order of micrometers (~20um - 200um), its length can be orders of magnitude greater (up to multiple meters, but generally upwards of a few centimeters). Embedding thus facilitates the manipulation (sectioning) of this material. However, many individuals have scalp hair that is not straight. To ensure a perpendicular cut, curled hair fibers have to be stretched while the resin hardens; a process that takes upwards of 24 hours. Alternatively, great care has to be taken to find the correct angle for a perpendicular cut after embedding. Regardless, cutting epoxy resin is an additional challenge, as it requires expensive, specialized equipment (such as a microtome). There are no widely used protocols that make it possible for the full morphological range of human scalp hairs to be embedded successfully.

**Methods for embedding & sectioning hairs are laborious.**

For the cross-sectional study of hair, the main hindrance is the need to embed the hair in resin. Especially for tightly coiled hairs, there is a major obstacle in having to find a way to keep the hair taut for the 24 hours the resin requires to harden. Attempts at embedding are not consistently successful (see Figure 1a). The main alternative has been to place the hair between glass slides and record width along the length of the fiber.(Hrdy, 1973; Trotter, 1930) But this is an inferior alternative because it does not allow for the visualization of the cross-section, it only gives us the longitudinal diameter of the hair.

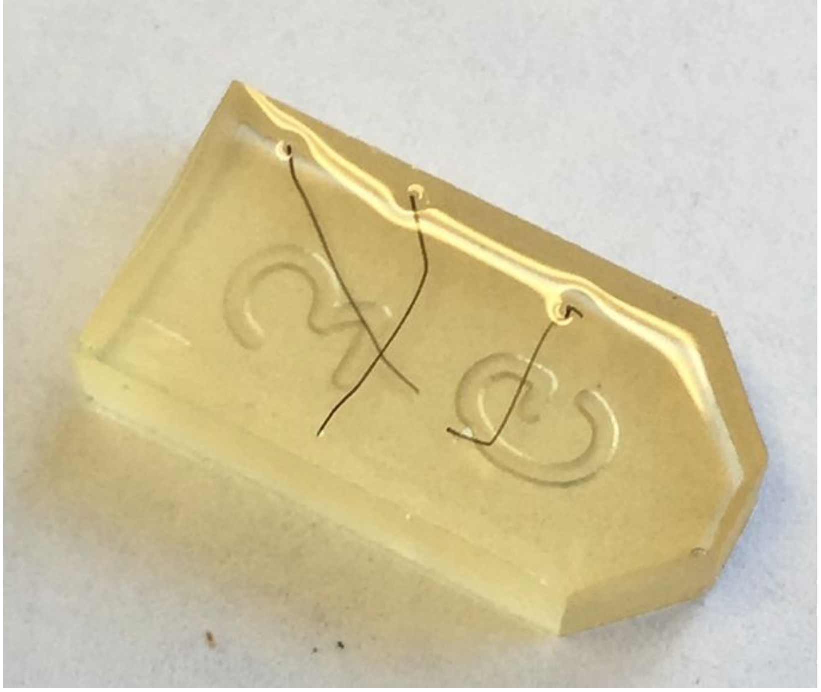

Figure 1a. Example of hairs moving during resin drying process

Some examples of existing protocols and methods for cross-sectional analysis include:

Trotter (1930) uses the longitudinal diameter as a proxy for the cross-sectional shape and size: “From each sample measurements were made on ten hairs chosen at random-thus a study of 3400 hairs from 340 individuals was made. Before measuring, the hairs were dipped into a solution of equal parts of ether and 95 per cent alcohol and then thoroughly dried; this process removed all dust and extraneous matter. Measurements of the greatest and least diameters of the hair shafts were obtained by means of an ocular micrometer in the microscope (magnification X 120) used in conjunction with a hair rotator, similar in construction to the one described by Danforth(2). From these measurements were computed the hair index and the area of the cross-section of the shaft.”

Hrdy (1973) similarly described multiple measures of the cross-sectional diameter which was used as a proxy for the cross-sectional geometry:
“(1) Average diameter. The average diameter was measured by placing the hair between glass slides and measuring with a micrometer-equipped microscope. The length of the hair was rapidly scanned and measured at many different places along the shaft, and the average value (in μm) recorded.”

Reis et al. (2020) use a method that involves bundling the hair before embedding and sectioning and show a step-by-step figure in their article:

“To overcome the flaws of other hair cross-section methods previously described,1, 3, 4 we used an epoxy embedding medium to maintain the hair strands intact and close together through the whole processing.

First, we prepared the embedding media—Agar 100 epoxy resin (Agar Scientific, Essex, UK)—using the fabricant recommended formulation to obtain hard blocks. We used a small drop to slightly embed each bundle of hair, keeping the hairs united and parallelly oriented. Each bundle was then left to rest for half an hour. This step is not mandatory but makes it easier to perform the next step. After this, we cut a small sample (about 0.5 cm in length) with the help of a scalpel and placed it in the molds, covering it with the rest of the embedding media. Polymerization at 60°C was then carried out for 24  hours (Figure 2). To finish, ultrathin sections (1250  nm) were cut using an ultramicrotome and stained with toluidine blue.”

Another example of more elaborate preparation of samples for study of the ultrastructure is seen in Koch et al. (2018):
“Each hair sample, consisting of approximately 20 terminal hairs, was tied together in a bundle to align the hairs longitudinally for embedding and cross-sectioning. The dehydration and fixation process typically employed for samples prepared for electron microscopy was not conducted prior to embedding as this process was found to dehydrate and potentially alter the structure of the cuticle during preliminary analyses. The hairs were air dried, embedded in Spurr’s resin, and placed in an oven at 60 degrees with desiccant to polymerize. Ultrathin sections (~70 nm thick) were cut perpendicular to the length of the hair shaft using a Leica ultramicrotome (Germany) and a Diatome Ultra diamond knife with a 35 degree blade. Sections were exposed to chloroform vapor to reduce section folding and potential deformation from impact with the cutting blade. Sections were collected onto Formvar supported slot grids, stained with a double staining procedure using lead citrate and uranyl acetate, and observed with a FEI (USA) Tecnai 1200 transmission electron microscope (TEM) with an accelerating voltage of 80 kV. Images of hair cross-sections were collected at 4200× magnification with a 20 percent overlap = in area. Montaging of the images was attempted; however, not all images aligned correctly.”

The methods we propose fall in between the efficient, but superficial measuring of longitudinal diameter and the meticulous but intensive, hand-washing of hairs and microtoming described in Reis et al. (2020) and Koch et al. (2018). Moreover, we provide more extensive video and still image guidance than we have found in the cross-sectional literature. This is to ensure that the protocol can be applied by any, regardless of previous experience or skill.

### Previous work on curvature

In contrast to cross-sections, the quantitative study of hairs longitudinal geometry has lagged behind. Longitudinal hair morphology ranges from a straight fiber, to a tightly coiled helix. A helix is a curve in 3-dimensional space that can be mathematically described by three parameters: arc length, curvature and torsion. The first attempt to operationalize the quantification of human scalp hair curvature appears relatively recently in the literature Hrdy's(1973) curvature quantification method requires a hair to be pressed between two glass slides to collapse three-dimensional variation into two dimensions. Once between the slides, a transparent template with circles of known radius are placed over the sample and matched to the curves of the hair. Despite the introduction of these methods, it has only been rarely applied to research on human scalp hair.(Bailey & Schliebe, 1986; Lasisi et al., 2016; Loussouarn et al., 2007)

#### Methods to quantify curvature have issues with replicability

Measuring curvature objective requires that the helical structure of non-straight hair fibers be reduced to two-dimensional curves. Previous curvature methods rely, without exception, on the subjective evaluation of the observer to one degree or another.(Lasisi et al., 2016; Loussouarn et al., 2007; Mkentane et al., 2017) Comparing hairs of different lengths is also a significant challenge. Depending on the method, a potential confounding factor is the greater number of points of measurement for hairs with high curvature as compared with low curvature (see Fig. 1b), involving potentially greater inter-observer variability in the number of and value of circles fitted to curves along a hair. Most importantly, of these existing methods, neither those for quantifying cross-section nor curvature could feasibly scale to the hundreds of samples many disciplines would need to phenotype.

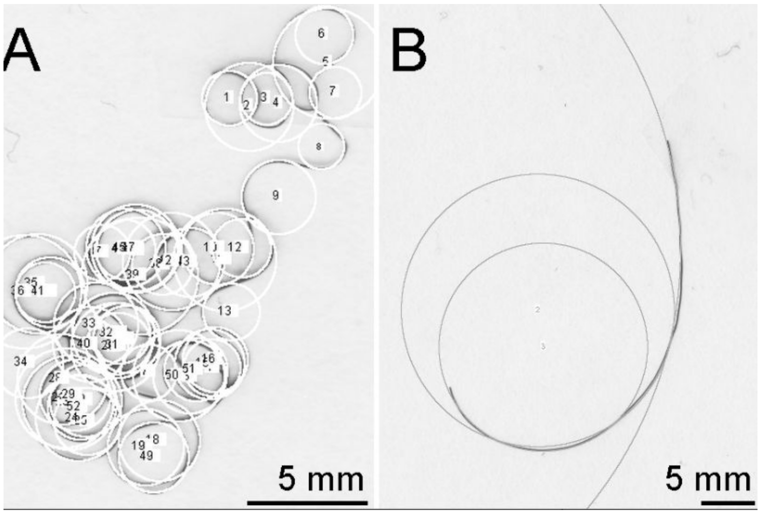


Figure 1b. Example of discrepancy in number of measurements taken in high vs. low curvature hair with previous methods.

Hrdy (1973) is the earliest published article describing a method of directly quantifying hair fiber curvature:

“(5) Average curvature. Each hair was placed between two glass slides, allowing measurement of the curvature of hairs that vary in three dimensions. The radius of curvature of each curve of the hair was determined by placing a transparent template with circles of known radius over the sample and matching an arc of the appropriate circle with the curve. Average curvature itself is the inverse of the average radius of curvature; a high average curvature is represented by a high number.”

Bailey and Schliebe (1986) is a test of precision for Hrdy’s original 1973 method:
“Since that time, this measurement has been found to be useful in the forensic comparison of curly, human head hairs with certain modifications made to the original method. The method currently used by this laboratory consists of the following steps:

1. Placing the hair in boiling water to remove grooming agents and relax the hair;
2. Removing excess water and allowing the hair to dry at room temperature;
3. Placing the hair between two glass plates to reduce the curvature to two dimensions; and
4. Measuring the resultant curves with a circle template of known radius.

The average curvature is calculated as the inverse of the average radius in millimeters times 100. This measurement then ranges from 0 to 100 mm with the curlier hairs having the higher value. Straight hairs have an average radius of 100 mm or more and an average curvature of 0 mm. To help establish the precision of this method, a single, curly, Caucasian head hair was measured 30 times by one examiner, and independently, 30 times by a second examiner. The data comparing the results of these measurements are shown in Table 1.
The major sources of variation in the measurement are as follows:

1. Amount of drying the hair received after boiling;
2. Determination of which dimension was reduced when placed between the glass plates;
3. Judgment of the number of curves to be measured; and
4. Judgment of which circle radius gives the best fit.”

Loussouarn et al. (2007) describe a method that is derived from Hrdy’s curvature method, but differing mainly in its decision to take only one measurement (the smallest) as representative of a sample’s curvature. Additionally, they describe a number of steps that measure various aspects of curl, but are likely redundant and covarying with curvature. Moreover the final partition into the eight curl types appears to be somewhat arbitrary:

“The method requires very simple materials: two glass plates, tape, a simplified CD meter, a curl meter, and a ruler. The CD meter includes the four cut-off values derived from the segmentation tree for the classification of types I–IV. The curl meter is made of a 0.98-cm-diameter circle, allows the segmentation of types V and VI vs. VII and VIII. The ruler helps to constrain the hair to 80% of its length in order to separate type V from VI and VII from VIII. More precisely, this simplified method can be described in three steps. Initially, the hair is carefully laid on a glass plate, without applying any mechanical stress, in order to allow it to maintain its natural shape. A second glass plate is gently placed onto the hair, carefully avoiding any side shifting or sliding. The first step is the evaluation of the curvature using the CD meter. The area of the CD meter where the smallest curvature is located indicates whether the hair is type I, II, III, or IV (Fig. 3). If the smallest curvature is included in the filled circle, the hair is type V, VI, VII, or VIII. Two other steps are needed to classify the hair. The second step is the test of curliness using the curl meter. The curl meter is placed on the glass plate in order to determine whether or not the hair fits entirely inside the circle (Fig. 4). The third and last step consists of counting the number of wave crests. The cover plate is removed. One end of the hair is taped in front of 0 cm on the ruler and the other end is taped at 4 cm. The two ends of the hair are taped on the bottom glass plate. Each self-stick strip covers 0.5 cm of hair fiber, and the distance between the two tapes is fixed to 4 cm thanks to the ruler. After replacing the cover plate, the hair takes a sigmoid form from which the highest number of wave crests is counted (Fig. 5). Based on the curl meter result, and on the number of wave crests, the hair type (V, VI, VII, or VIII) can be defined with the following rules. If the entire hair is included in the curl meter and the number of crests is from one to five, the hair type is VII. If the entire hair is included in the curl meter and the number of crests is six or more, the hair type is VIII. If the entire hair is not included in the curl meter and the number of crests is from one to three, the hair type is V. If the entire hair is not included in the curl meter and the number of crests is four or more, the hair type is VI."

Lastly, Lasisi et al. (2016) propose something that can best be described as a direct digital application of Hrdy's original curvature method:
"Single strands of hair were placed on a sheet of blank white paper, the application of mechanical stress was avoided to ensure unaltered curvature, and the paper was covered with a transparent sheet of acetate to facilitate a two-dimensional measurement. Three hairs were analyzed per person in order to provide a representative average for each individual. Samples were scanned using a flatbed scanner (CanoScan LiDE 600F) at a resolution of 1200 DPI. Curvature measurements were collected in ImageJ version 1.48. Each curve in a hair fiber was traced digitally, creating a separate data point from each curve. Curvature varied greatly among individuals, thus the number of measurements per hair varied from as few as two to as many as 60 (Fig. 1). The mean, maximum, and minimum curve diameters from a single hair are then computed. Curvature is the inverse of the radius (curve diameter ÷ 2), so the previously computed values yield a mean, maximum, and minimum curvature. The variable average curvature is the mean curvature averaged from the three hairs analyzed for a single individual. The variable irregularity is similarly calculated as an individual's averaged maximum to minimum curvature ratio. Both average curvature and irregularity variables are derived from Hrdy (1973). Precise instructions for the method of curvature analysis described in this study have been made available (Supporting Information Document 2) and a depiction of the application of this digital method can be seen in Figure 1."

While Lasisi et al. 2016 presented an extensively documented methodology, this protocol was very laborious and did not include a washing step that was scalable (
