## Supplementary Results 1 for "High-throughput phenotyping methods for quantifying hair fiber morphology"

### Validation

Tina Lasisi

2020-11-23 22:09:24

#### Curvature

Here, we will evaluate the accuracy of *fibermorph* in estimating the length and curvature of hair using simulated data. See simulation script here.

The simulated data can be found here.

We simulated arcs of various curvatures at a length of 1.57mm. There were 25 arcs per image.

##### Simulated vs. estimated curvature & length

To calculate the accuracy of our measurements, we compared the known parameters with the parameters estimated from our fibermorph package.

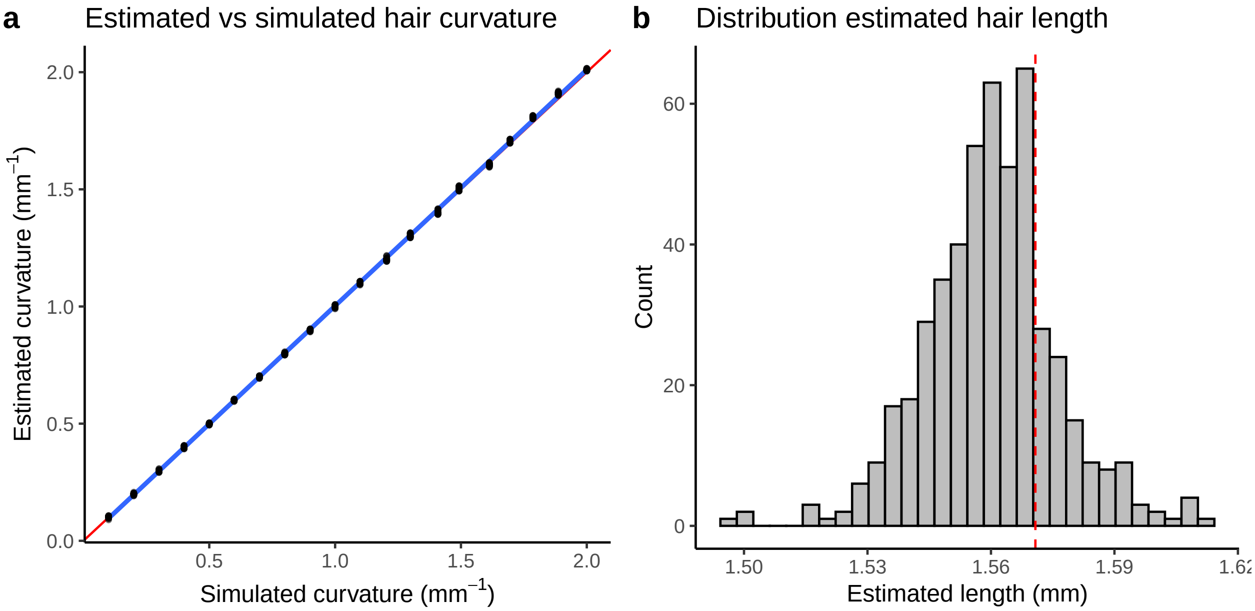


In Fig.a we see that there is a near perfect correlation between the simulated and estimated curvatures. Fig.b shows the distribution of estimated hair lengths around the simulated length (red line).

We plot simulated curvature against estimated length to show the distribution of estimated length as a function of curvature.

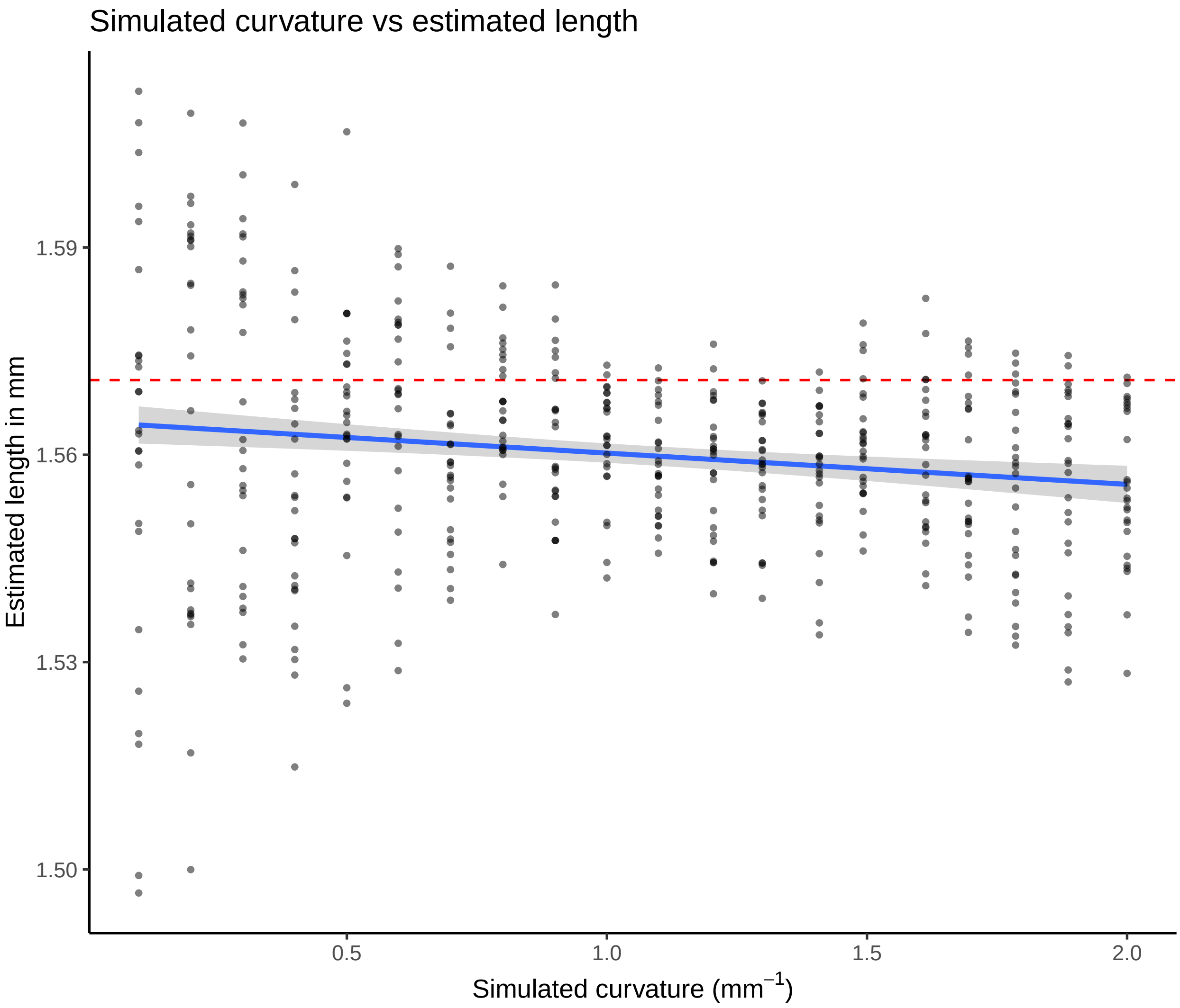


There seems to be a broader range of error in the estimation of length in straighter hairs. This is likely a result of the majority of pixels being oriented in a manner that causes a divergence between the pixel length (number of pixels) and the real length that is being measured. We apply a correction for this known issue in image analysis, however, it is expected that there will still be some error. Note that each point in this figure represents an individual hair fragment within an image. This supports the notion that it is not the low curvature per se, but rather the combination of low curvature and specific orientations that increases the error in length estimation.

#### Measurement error in curvature and length

In addition to the correlations between simulated and estimated parameters, we calculate root mean square error (RMSE) and percent error as alternatives to investigate the measurement error of our package.

NB: we present the data summarized for each image (i.e. all 25 fragments) as we cannot provide a hair fragment to hair fragment comparison.

##### Error statistics

Below, we calculate the mean error values for both RMSE and percent error.

Table 1: RMSE and Percent Error per variable

| var | mean.rmse | perent.error |
|---|---|---|
| curvature | 0.0002210 | 0.4720430 |
| length | 0.0004312 | 0.6863358 |
| radius | 0.0004624 | 0.4626120 |

We see less than 1% error across the variables and RMSE of less than 0.0005.

Below, we plot the data.

##### Root mean square error

First, we plot the root mean square error for curvature and length.

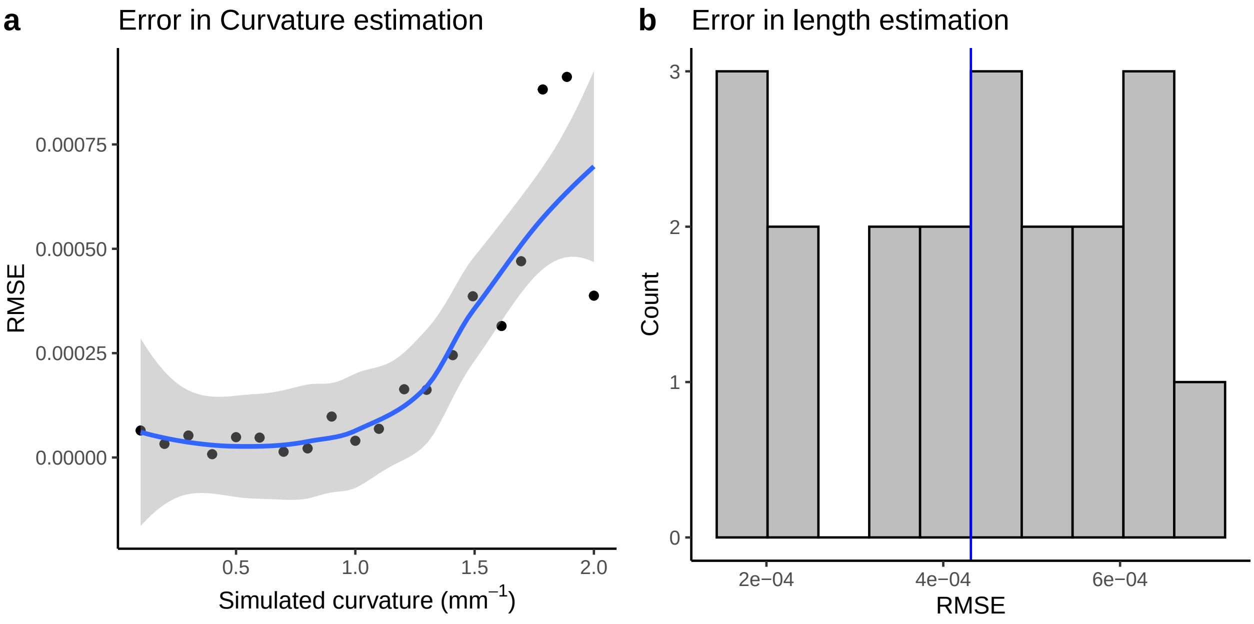


We then examine the relationship between curvature and RMSE of length

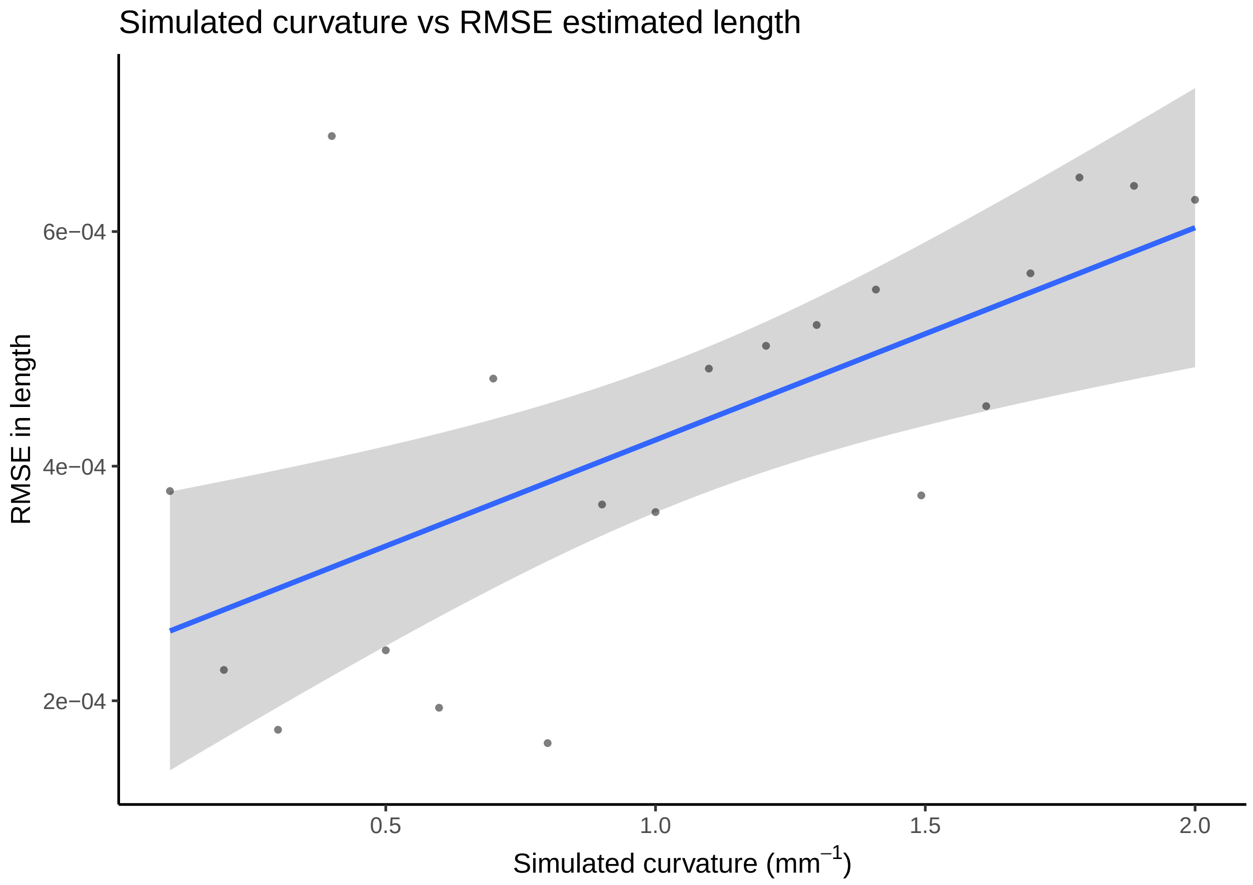


We observe an increase in RMSE with curvature.

##### Percent error

Below we plot the percent error for curvature and length.

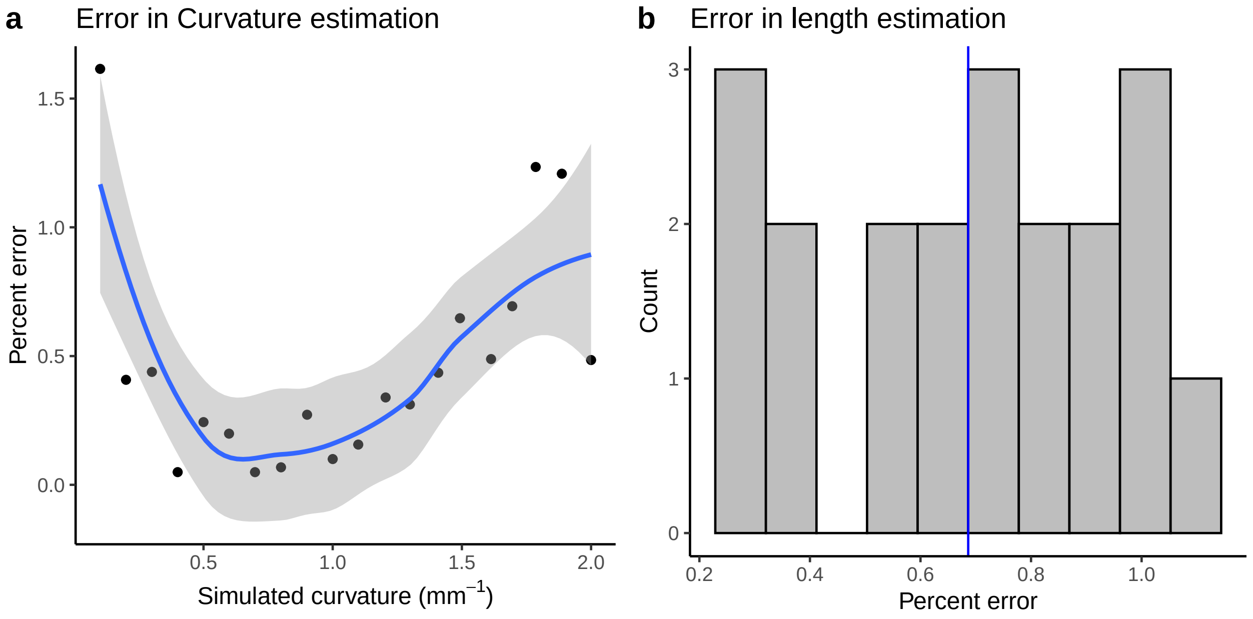

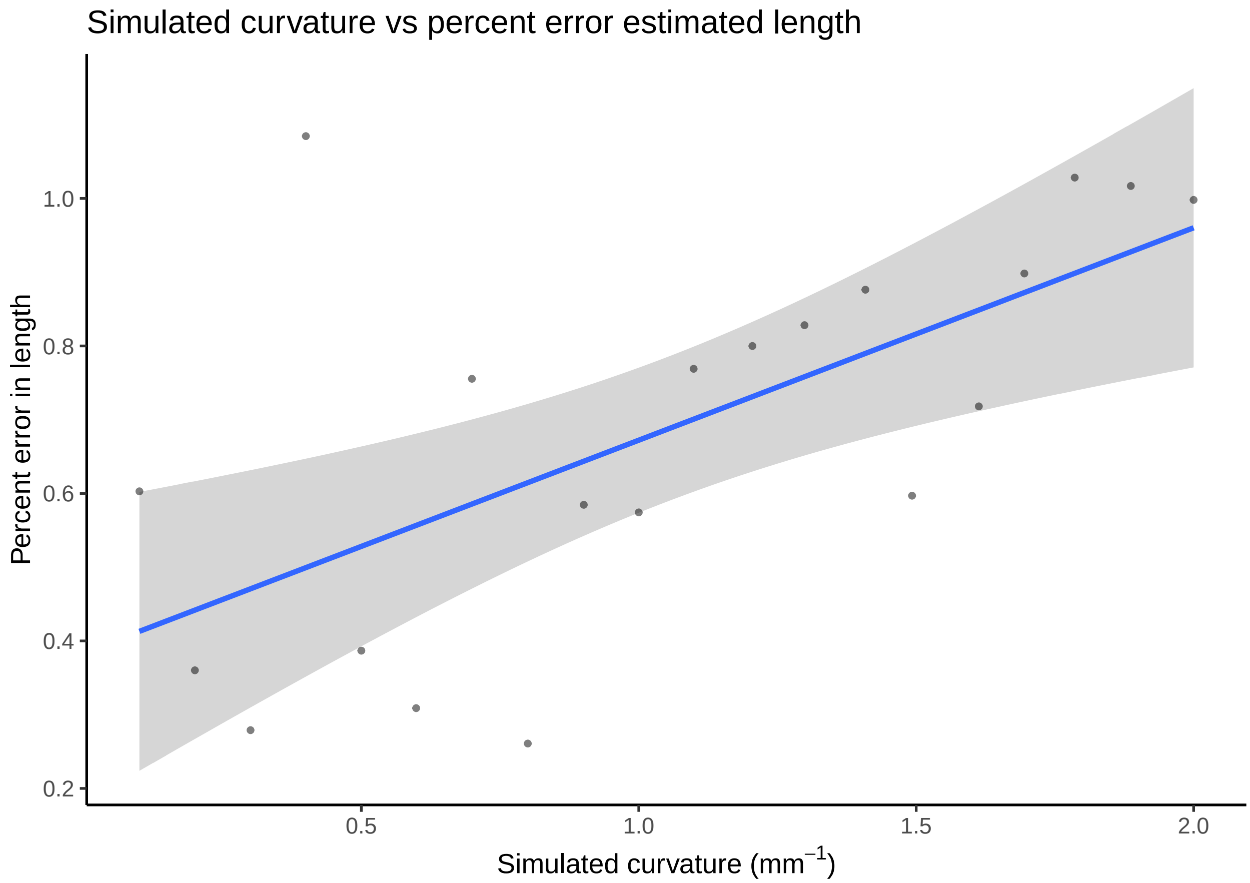


Here we see that error appears to increase slightly with curvature if considering the data in terms of percent error.

#### Cross-section

The fibermorph section function estimates area, minimum diameter, maximum diameter and eccentricity for a given cross-sectional image. We tested the measurement error using randomly generated circles and non-circular ellipses.

#### Correlation between simulated and estimated section parameters

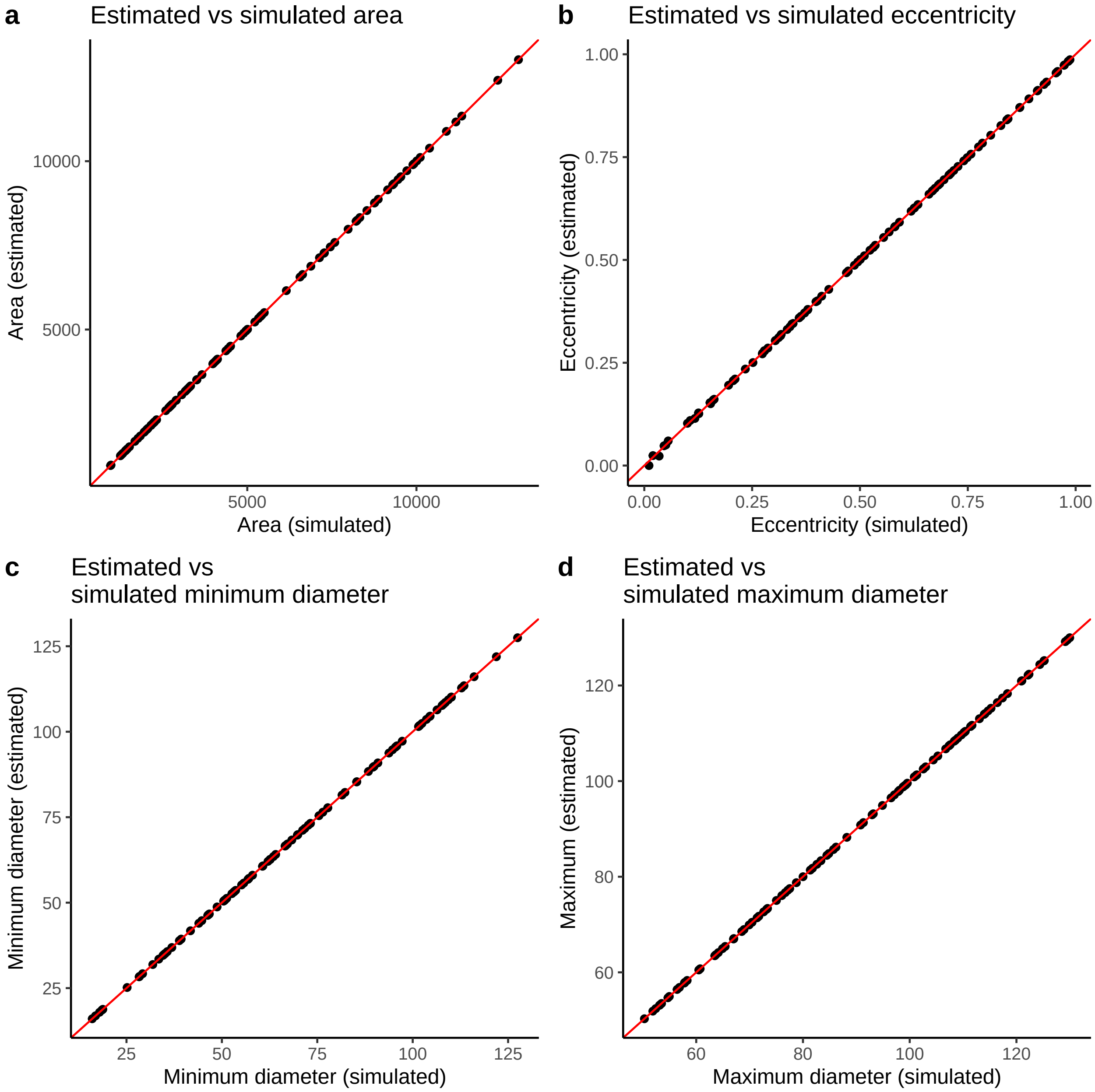


We see strong correlations between the estimated and simulated values for each cross-sectional parameter.

#### Measurement error for cross-sectional parameters

We calculate the percent error and RMSE for the cross-sectional parameters.

First, we calculate mean error values for all parameters.

Table 2: RMSE and Percent Error per variable

| var | mean_rmse | perent_error |
|---|---:|---:|
| area | 0.5136320 | 0.0137703 |
| eccentricity | 0.0007514 | Inf |

| var | mean_rmse | perent_error |
|---|---|---|
| max | 0.0097800 | 0.0120605 |
| min | 0.0080884 | 0.0136924 |

Percent error is considerably under 0.02% for each of the parameters with RMSE under 0.01 for all but area.

As one of the simulated ellipses was a circle with an eccentricity of 0, any deviation from this produces an infinite percent error. So below we present the values removing this observation.

Table 3: RMSE and Percent Error per variable

| var | mean_rmse | perent_error |
|---|---|---|
| area | 0.5136320 | 0.0137703 |
| eccentricity | 0.0006492 | 1.0066337 |
| max | 0.0097800 | 0.0120605 |
| min | 0.0080884 | 0.0136924 |

##### Root mean square error

Below, we plot RMSE as a function of each parameter.

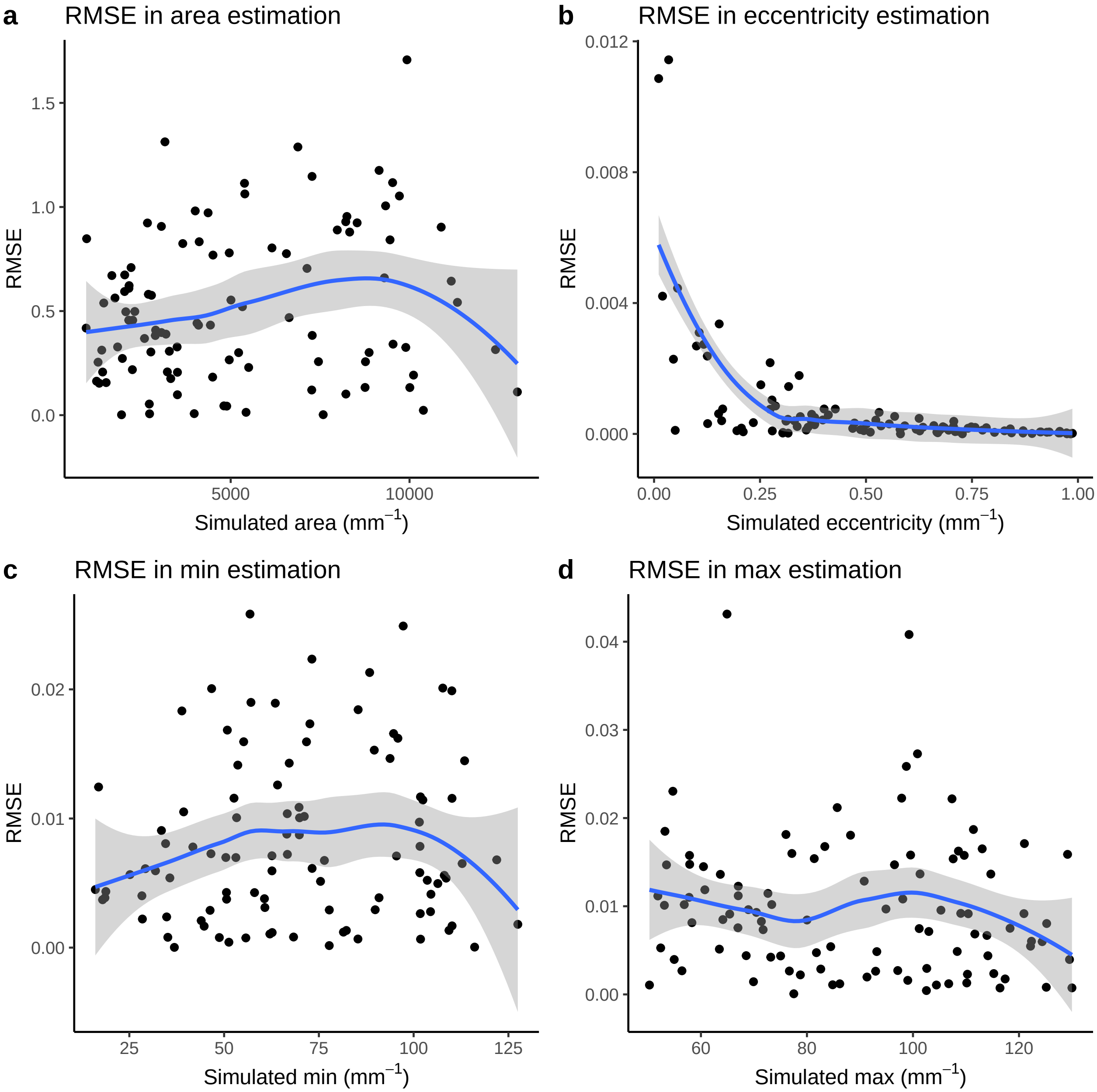


There does not appear to be any overarching pattern in RMSE across the variables.

##### Percent error

Below we plot the correlation between simulated values and percent error for each parameter.

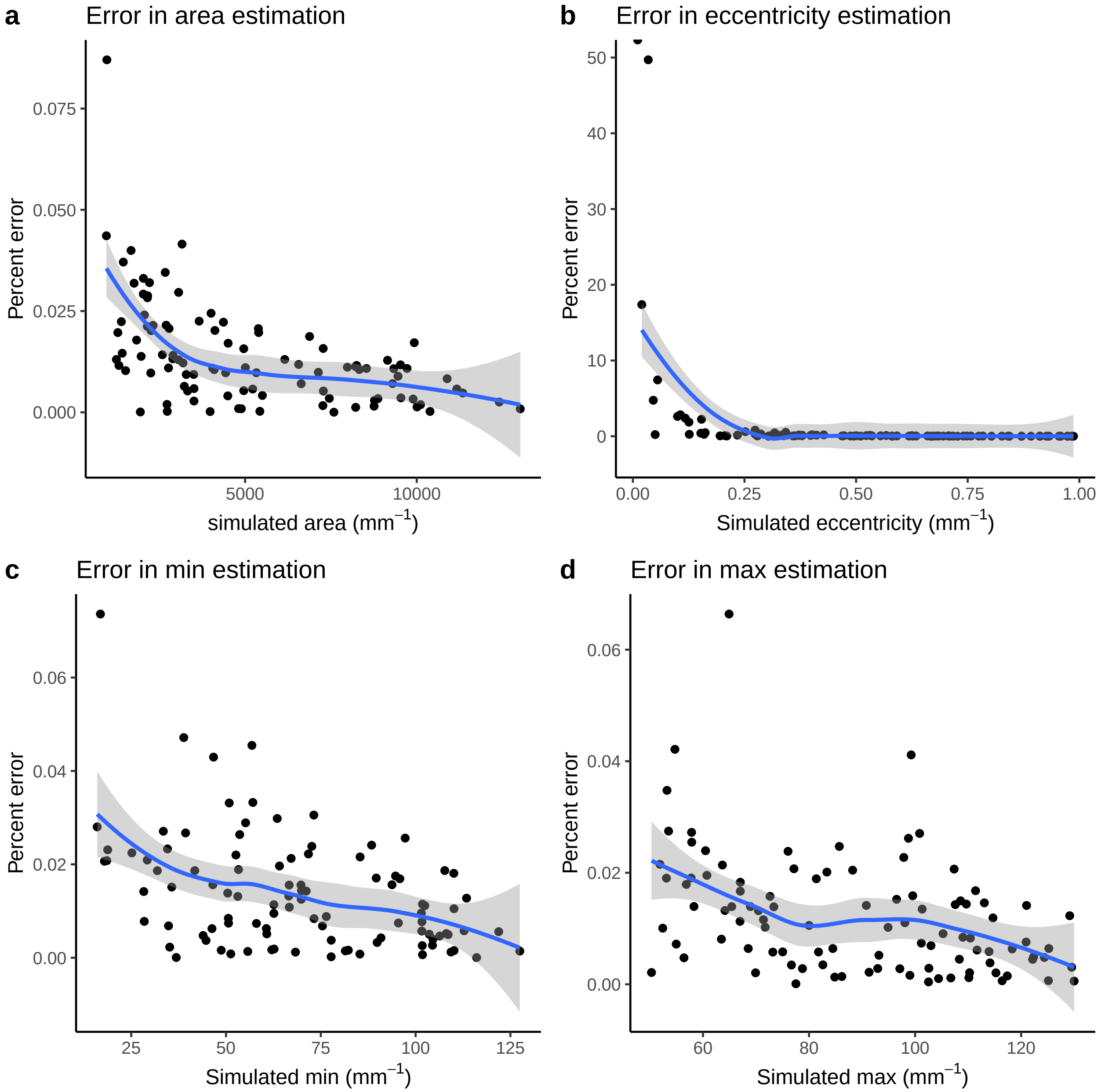


We observe a general decrease in percent error for each parameter.
