## Supplementary Results 2 for "High-throughput phenotyping methods for quantifying hair fiber morphology"

### Biological Significance

Tina Lasisi

2020-11-24 15:29:16

To explore the significance of quantifying hair fiber morphology, we explore the relationship between various quantitative hair traits, categorical data and genotype data on the same sample.

Our data consists of 193 individuals for whom we have quantitative hair phenotype data. In our first data quality control step, we filter to keep individuals who have more than 4 hair fragments in their curvature image and over 10% African ancestry. We calculate mean and median values for the cross-sectional data we have collected for individuals (~ 6 sectioned hair fibers). In our analyses, we use median values as they are less affected by intra-individual outliers.

#### Self-reported hair texture vs. quantitative

We compare the self-reported hair texture with mean and median curvature for our sample.

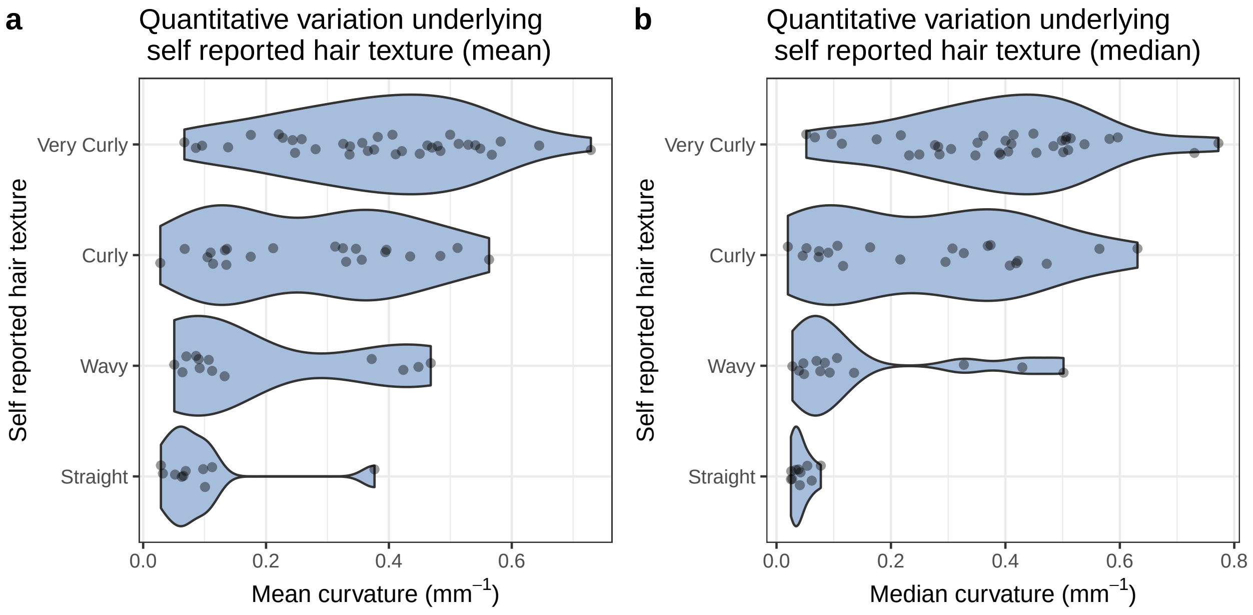


The single individual with a high mean curvature in the straight group is the result of an artefact in the image.

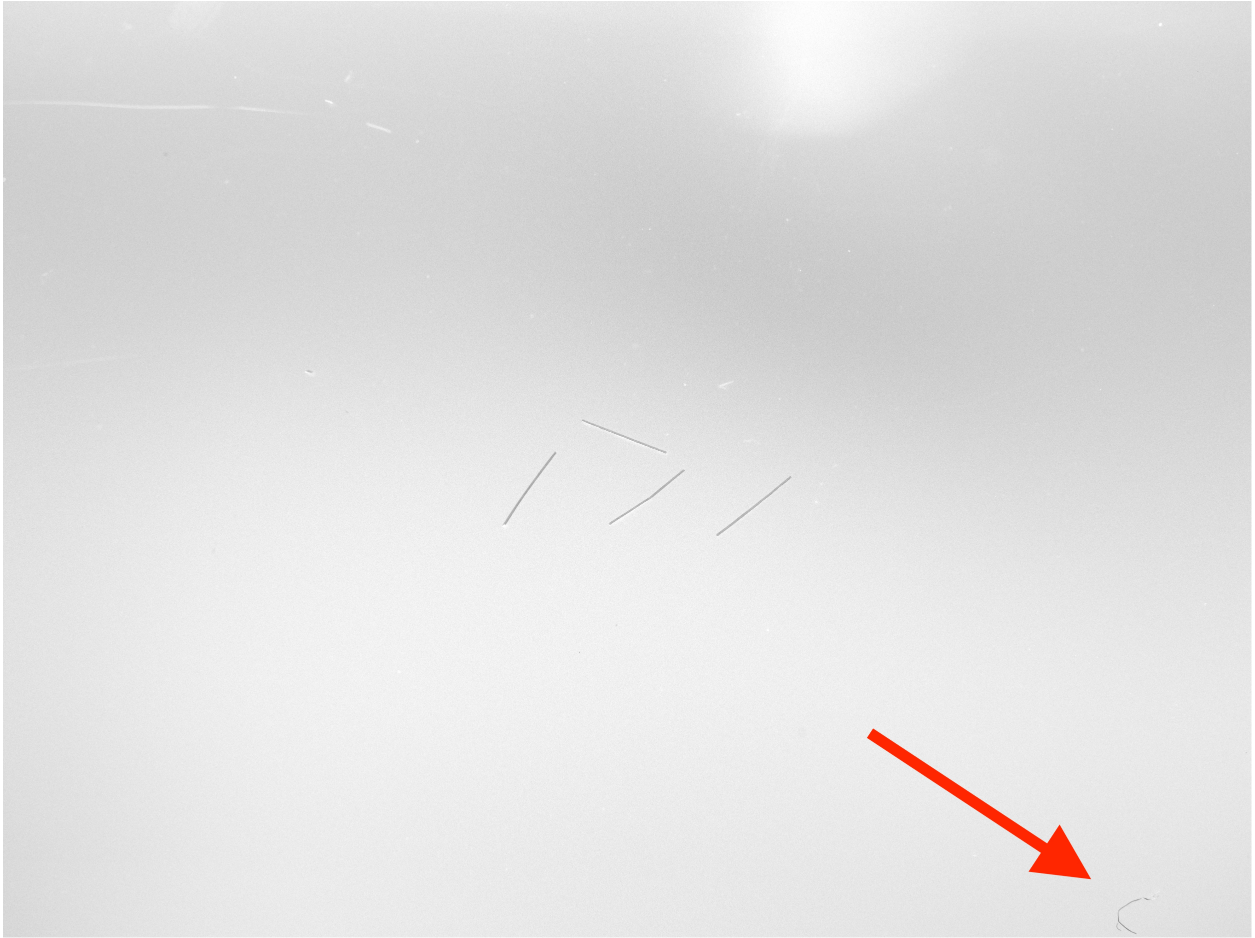

The red arrow points to a stray fiber that likely contaminated the sample and was missed during imaging. Such potential outliers are the reason we chose to use the median curvature for a sample in our analyses.

### Objective hair texture vs. quantitative

To explore how much data is lost when binning continuous variation, we compared mean and median curvature to classified hair texture. This classification is based on Loussouarn et al.'s 2007 paper "Worldwide diversity of hair curliness: a new method of assessment.

While the authors propose a number of parameters to distinguish curlier hair types (based on number of twists and waves among other factors), their primary classification is based on curvature. We demonstrate that, regardless of additional parameters, a considerable range of curvature is obscured when collapsing hair variation according to their curvature thresholds.

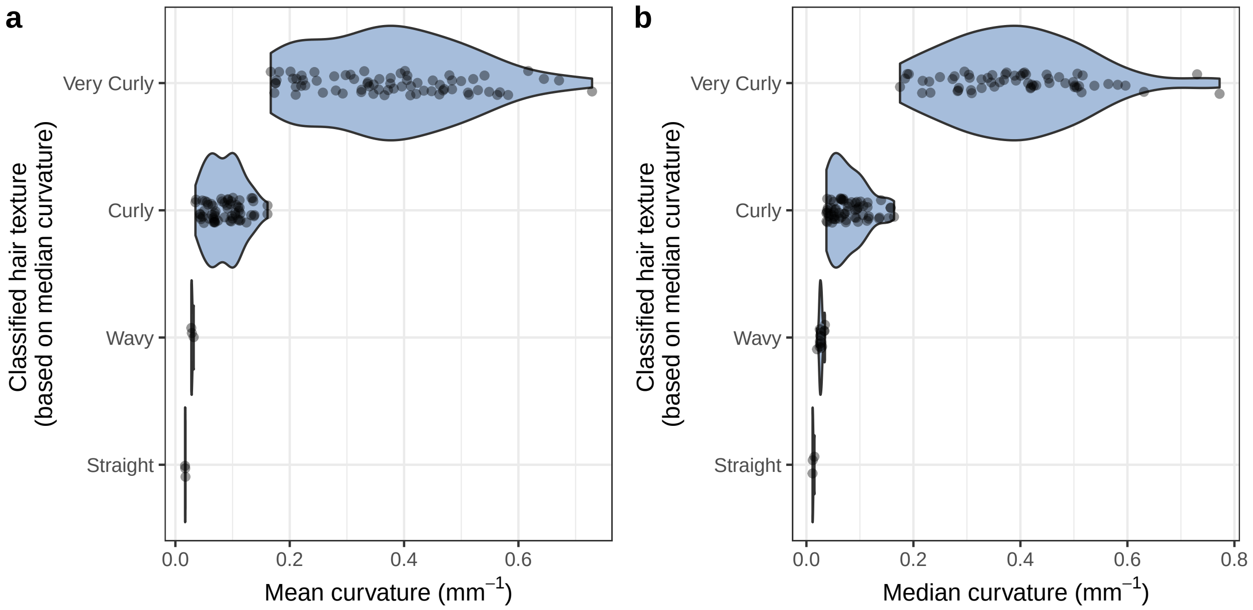


#### Ancestry vs. hair morphology

We carried out a number of analyses using the genotype data collected for this diverse sample. In an admixed sample where a continuous trait has divergent distributions in the parental ancestry groups, the resulting admixed population can show a correlation between ancestry and that trait. Finding such a correlation suggests may imply a polygenic trait with high heritability.

##### Admixture components

Our sample consists of admixed individuals with primarily African and European ancestry.

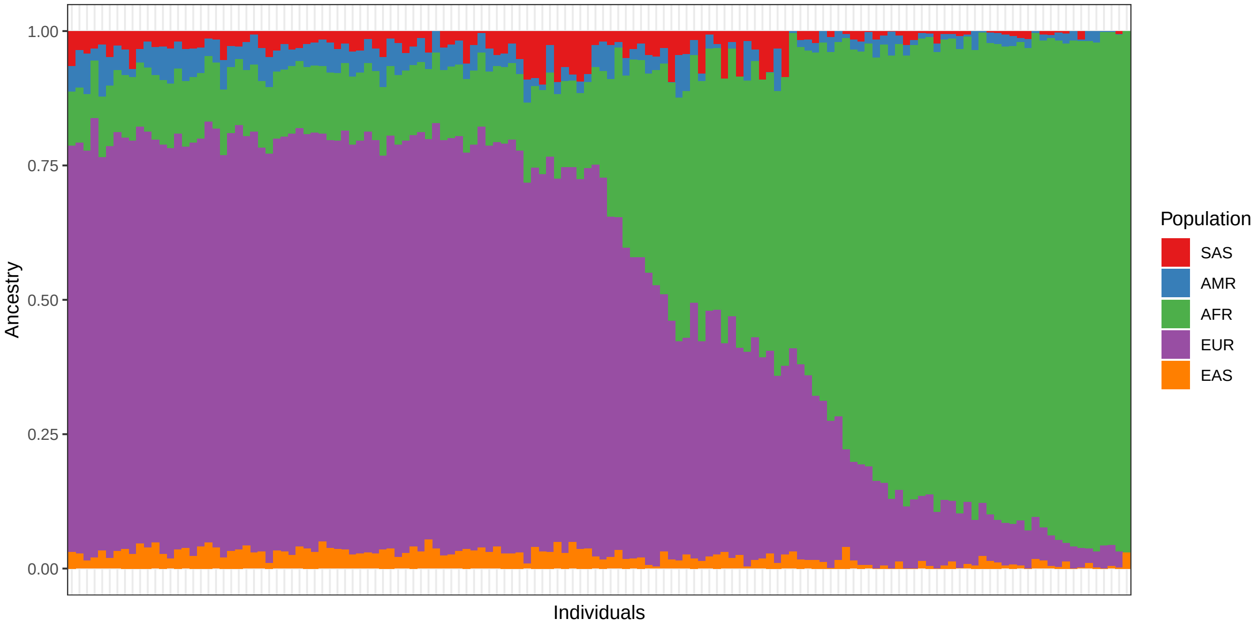


The colors represent ancestries that correspond to the following 1000 Genomes populations: - SAS = South Asian - AMR = American - AFR = African - EUR = European - EAS = East Asian

Each of these are metapopulations based on the grouping of multiple (sub)continental population groups in the 1000 Genomes repository.

#### Ancestry vs. curvature

Here we plot the correlation between proportion of African ancestry and m-index, median curvature, and eccentricity.

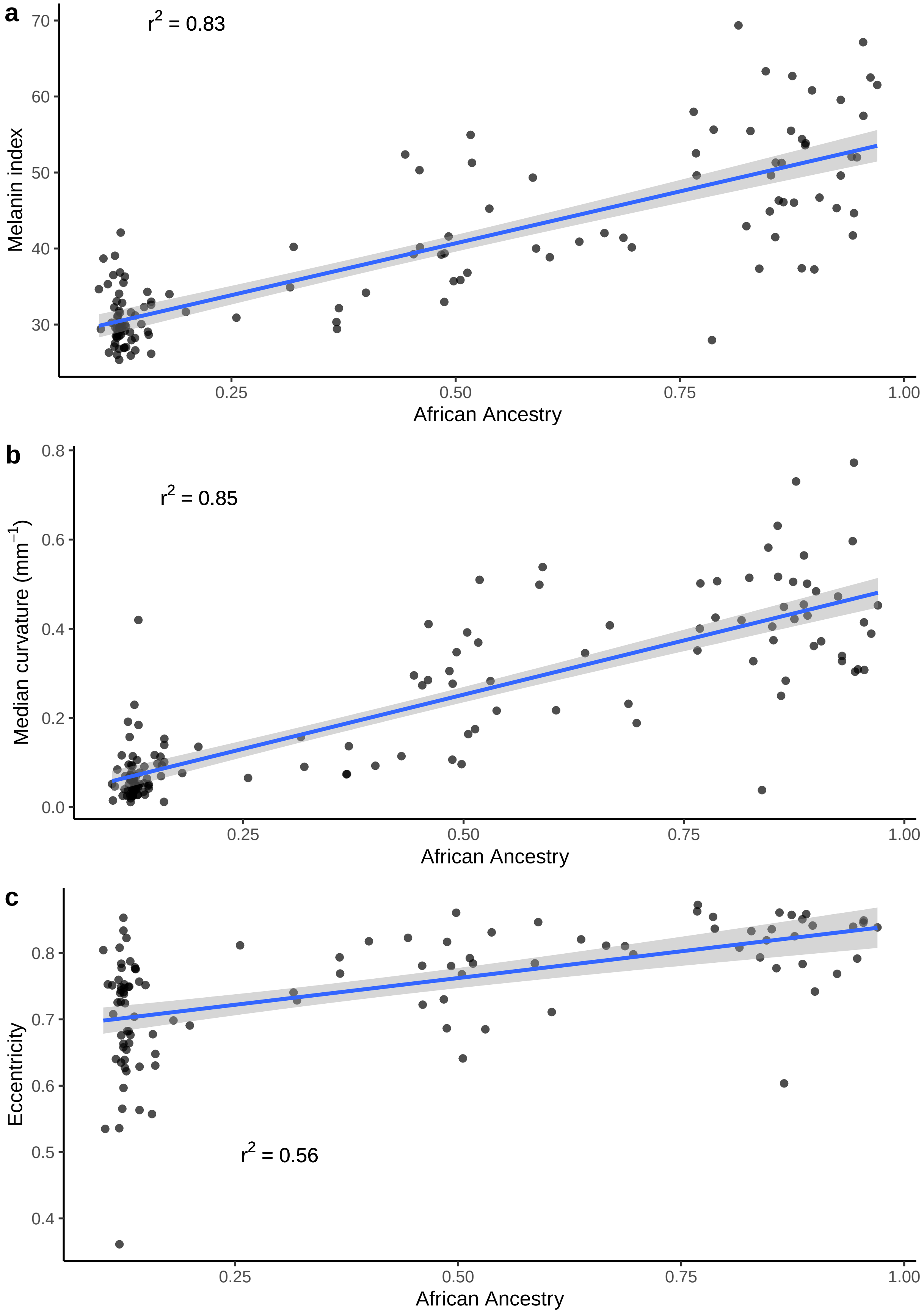
a
r2 = 0.83
Melanin index
70
60
50
40
30
0.25
0.50
0.75
1.00
African Ancestry
b
r2 = 0.85
Median curvature (mm−1)
0.8
0.6
0.4
0.2
0.0
0.25
0.50
0.75
1.00
African Ancestry
c
r2 = 0.56
Eccentricity
0.8
0.7
0.6
0.5
0.4
0.25
0.50
0.75
1.00
African Ancestry

#### Curvature vs. eccentricity

The relationship between cross-sectional shape (eccentricity) and curvature has long been debated. Due to the coincidence of cross-sectional shape and curvature in various populations that are often contrasted (i.e. East Asian vs. North European vs. West African), it has been unclear whether these traits have a causal relationship (specifically that higher eccentricity predicts higher curvature). In our admixed sample, we have the opportunity to test this question and fit a model between these traits with and without ancestry.

##### Uncorrected

First we examine the data without correcting for ancestry.

```
##
## Call:
## lm(formula = curv_median ~ eccentricity_median, data = df_curv_ecc)
##
## Residuals:
##     Min      1Q  Median      3Q     Max
## -0.2843 -0.1245 -0.0264  0.1125  0.4770
##
## Coefficients:
##                     Estimate Std. Error t value Pr(>|t|)
## (Intercept)          -0.5682     0.1270  -4.474 1.94e-05 ***
## eccentricity_median   1.0284     0.1692   6.076 1.96e-08 ***
## ---
## Signif. codes:  0 '***' 0.001 '**' 0.01 '*' 0.05 '.' 0.1 ' ' 1
##
## Residual standard error: 0.1576 on 106 degrees of freedom
## Multiple R-squared:  0.2583, Adjusted R-squared:  0.2513
## F-statistic: 36.92 on 1 and 106 DF,  p-value: 1.959e-08
```

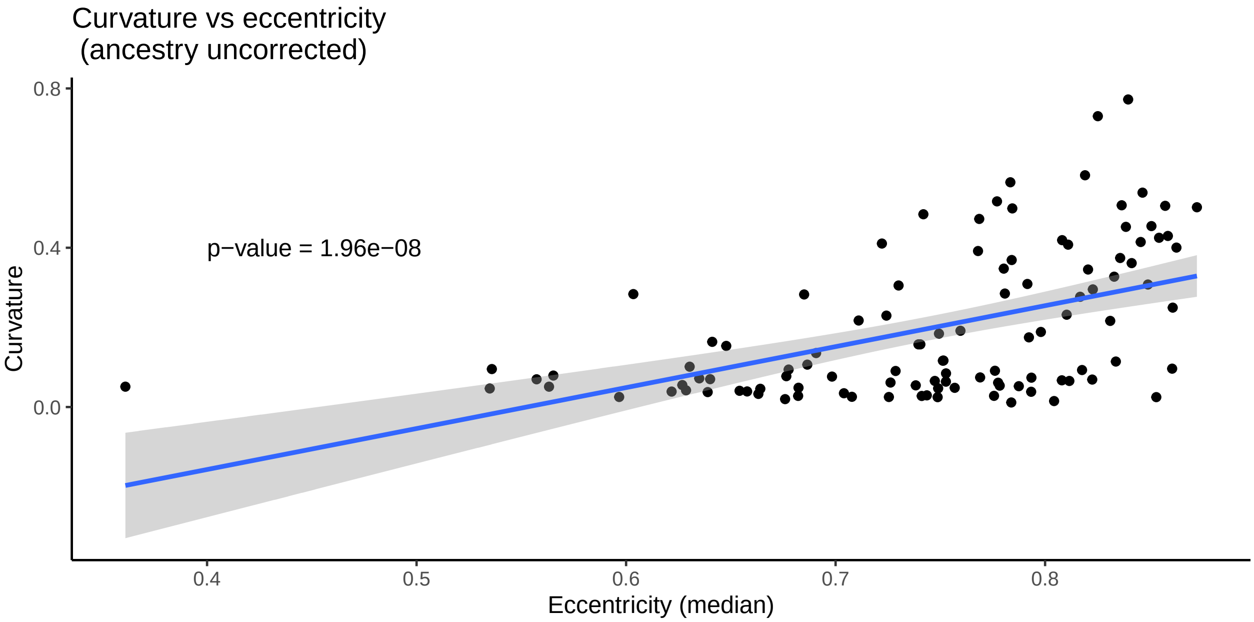


If we consider the relationship between curvature and eccentricity without taking into account ancestry, we find that eccentricity is a significant predictor of curvature.

##### Corrected

We then re-analyze the data with ancestry as a covariate.

```
##
## Call:
## lm(formula = curv_median ~ eccentricity_median + AFR, data = df_curv_ecc)
##
## Residuals:
##      Min       1Q   Median       3Q      Max
## -0.37902 -0.04122 -0.00657  0.03606  0.29971
##
## Coefficients:
##                     Estimate Std. Error t value Pr(>|t|)
## (Intercept)         -0.07237    0.08661  -0.836    0.405
## eccentricity_median  0.10891    0.12511   0.871    0.386
## AFR                  0.48101    0.03632  13.243   <2e-16 ***
## ---
## Signif. codes:  0 '***' 0.001 '**' 0.01 '*' 0.05 '.' 0.1 ' ' 1
##
## Residual standard error: 0.09693 on 105 degrees of freedom
## Multiple R-squared:  0.7222, Adjusted R-squared:  0.7169
## F-statistic: 136.5 on 2 and 105 DF,  p-value: < 2.2e-16
```

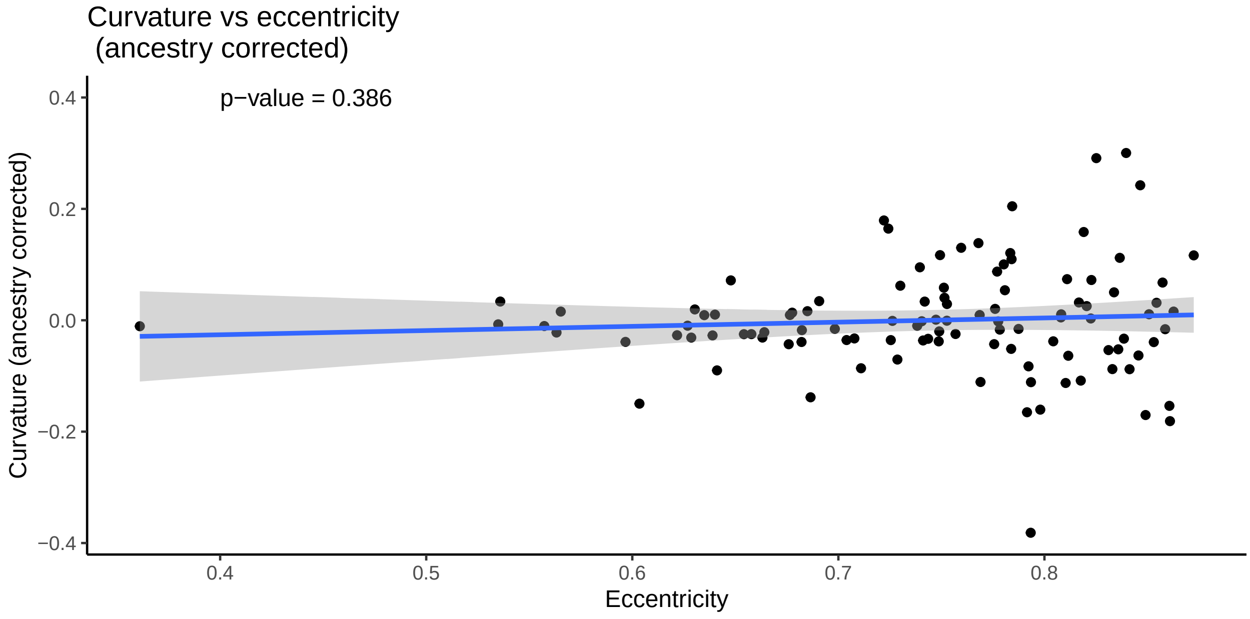


However, when we correct for ancestry, this correlation is no longer significant. This supports the idea that these traits may be independent.

#### Curvature vs. skin pigmentation

To demonstrate the potential effect of population stratification on traits, we compare hair curvature with skin pigmentation (m-index). These two traits are not biologically related, yet, in an admixed population, we may see a correlation that is due to population stratification of these polygenic traits.

##### Uncorrected

First we examine the relationship between curvature and skin pigmentation without correcting for ancestry.

```
##
## Call:
## lm(formula = curv_median ~ m_index, data = df_curv_mindex)
##
## Residuals:
##      Min       1Q   Median       3Q      Max
## -0.20715 -0.06804 -0.03471  0.04818  0.51483
##
## Coefficients:
##              Estimate Std. Error t value Pr(>|t|)
## (Intercept) -0.280232   0.041533  -6.747 4.91e-10 ***
## m_index      0.012887   0.001031  12.501  < 2e-16 ***
## ---
## Signif. codes:  0 '***' 0.001 '**' 0.01 '*' 0.05 '.' 0.1 ' ' 1
##
## Residual standard error: 0.1261 on 126 degrees of freedom
## Multiple R-squared:  0.5536, Adjusted R-squared:  0.5501
## F-statistic: 156.3 on 1 and 126 DF,  p-value: < 2.2e-16
```

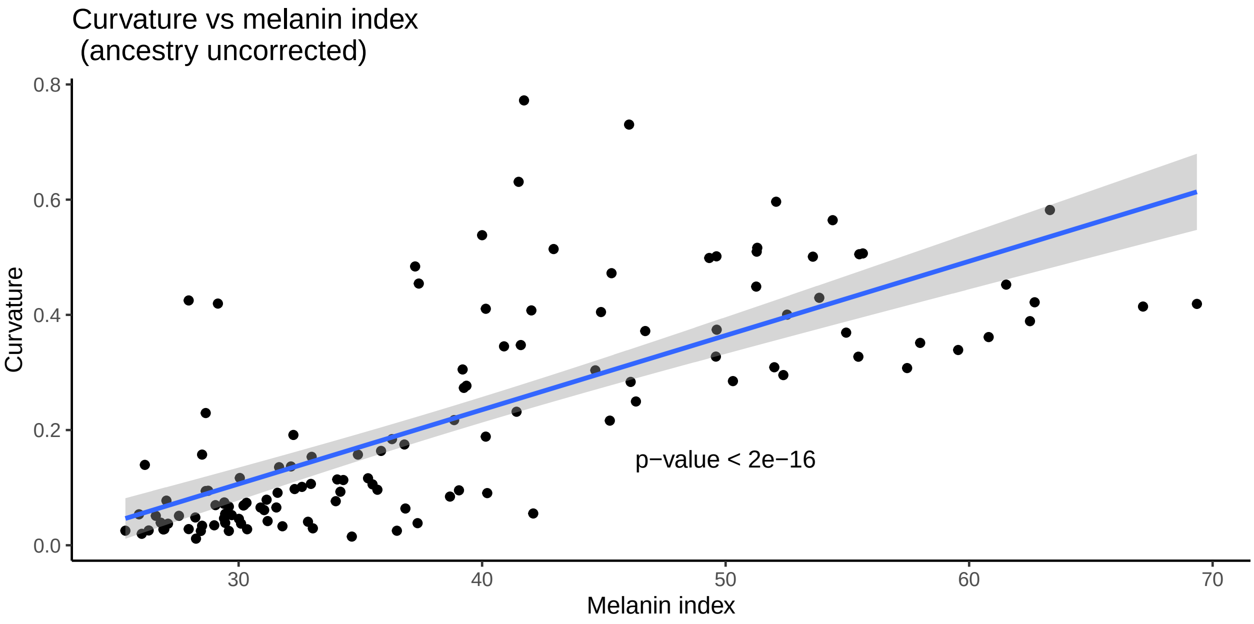


As expected, we see a significant correlation between the two traits.

##### Corrected

We then apply a correction for ancestry and re-analyze the data.

```
##
## Call:
## lm(formula = curv_median ~ m_index + AFR, data = df_curv_mindex)
##
## Residuals:
##      Min       1Q   Median       3Q      Max
## -0.34321 -0.04877 -0.01348  0.03562  0.34806
##
## Coefficients:
##              Estimate Std. Error t value Pr(>|t|)
```

```
## (Intercept) -0.061501   0.042384  -1.451    0.149
## m_index      0.002734   0.001469   1.862    0.065 .
## AFR          0.406609   0.048561   8.373 9.63e-14 ***
## ---
## Signif. codes:  0 '***' 0.001 '**' 0.01 '*' 0.05 '.' 0.1 ' ' 1
##
## Residual standard error: 0.1014 on 125 degrees of freedom
## Multiple R-squared:  0.714,  Adjusted R-squared:  0.7094
## F-statistic:   156 on 2 and 125 DF,  p-value: < 2.2e-16
```

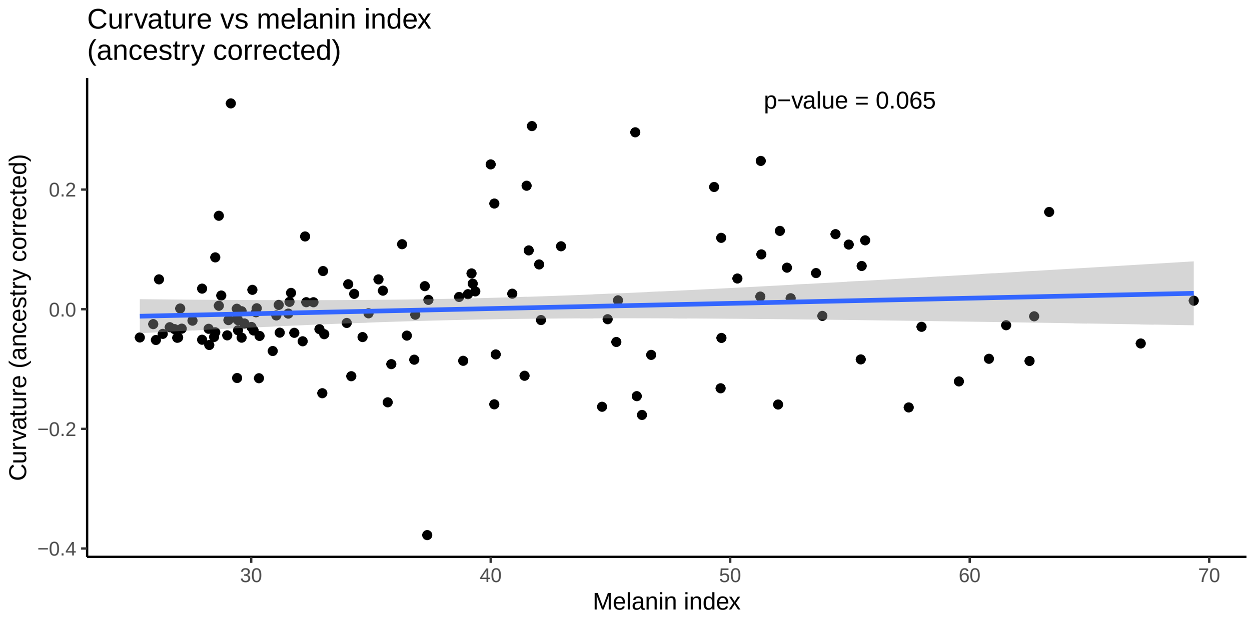


Like with curvature and eccentricity, the relationship between curvature and skin pigmentation is no longer significant when ancestry is taken into account.
